# Computational modeling reveals a key role for polarized myeloid cells in controlling osteoclast activity during bone injury repair

**DOI:** 10.1101/2020.10.13.338335

**Authors:** Chen Hao Lo, Etienne Baratchart, David Basanta, Conor C Lynch

## Abstract

Bone-forming osteoblasts and -resorbing osteoclasts control bone injury repair, and myeloid-derived cells such as monocytes and macrophages are known to influence their behavior. However, precisely how these multiple cell types coordinate and regulate each other over time to repair injured bone is difficult to dissect using biological approaches. Conversely, mathematical modeling lends itself well to this challenge. Therefore, we generated an ordinary differential equation (ODE) model powered by experimental data (osteoblast, osteoclast, bone volume, pro- and anti-inflammatory myeloid cells) obtained from intra-tibially injured mice. Initial ODE results using only osteoblast/osteoclast populations demonstrated that bone homeostasis could not be recovered after injury, but this issue was resolved upon integration of pro- and anti-inflammatory myeloid population dynamics. Surprisingly, the ODE revealed temporal disconnects between the peak of total bone mineralization/resorption, and osteoblast/osteoclast numbers. Specifically, the model indicated that osteoclast activity must vary greatly (>17-fold) to return the bone volume to baseline after injury and suggest that osteoblast/osteoclast number alone is insufficient to predict bone the trajectory of bone repair. Importantly, the values of osteoclast activity fall within those published previously. These data underscore the value of mathematical modeling approaches to understand and reveal new insights into complex biological processes.

## INTRODUCTION

Bone healing subsequent to injury or trauma is a significant clinical problem in orthopedics and rehabilitation^1–3^. Understanding the processes involved and how cells coordinate and control each phase of injury repair can reveal opportunities to accelerate healing and improve patient outcomes while reducing cost. Currently, the phases of bone injury repair in diaphyseal, epiphyseal or metaphyseal fractures have been well characterized^1, 4–7^. For example, in critical non-union fractures, a rapid inflammatory response is followed by callus formation. The callus is then mineralized by infiltrating mesenchymal stromal cells (MSCs) that differentiate into cartilage, and bone-forming chondrocytes and osteoblasts respectively^1,2^. Subsequently, activated osteoclasts mediate resorption and clearing of the ossified callus^1^. In addition to osteoblasts and osteoclasts, other cell types are also involved in the bone healing process, such as resident and infiltrating immune cells that exert pro- and anti-inflammatory activities depending on environmental cues^1, 8–10^. This is evidenced by the fact that acute pro-inflammatory factor administration (e.g TNFα) can improve bone repair while prolonged administration has the opposite effect^11–15^. Monocytes and macrophages are major components of the bone immune infiltrate subsequent to injury^1, 8, 9, 16^. Previous studies using genetic or pharmacological depletion of myeloid cells such as macrophages demonstrated significantly delayed time to bone repair^10, 16–19^. However, precisely how these multiple cell types coordinate and regulate osteoblast and osteoclast activity over time is challenging to dissect using traditional *in vitro* and *in vivo* biological approaches.

A potential approach to overcome this hurdle is the integration of experimental data with computational models that allow for the analysis of multiple cells types at any time point during bone injury repair. Previous reports, including from our group, have successfully demonstrated the feasibility of mathematical modeling approaches to enhance our understanding of how cells interact in the bone ecosystem to coordinate homeostasis and cancer-bone interactions^20–29^. There are a number of mathematical model approaches that can be employed such as ODEs that can be used to model bone cell populations in normal and disease processes^30–35^. Individual cellular dynamics can also be considered by representing the cell populations as either a continuous spatial field whose dynamics are described by a set of partial differential equations (PDE)^36, 37^, or as individual agents in an agent-based model approach^30^. Although these models have been used to examine bone injury repair and homeostasis, they have largely focused on the interaction between bone-building osteoblasts and bone-resorbing osteoclasts^30–32, 34, 37^. Some models have considered immune populations but these are theoretical and are not driven by biological data that provides quantitative information for each population and various timepoints throughout the bone injury repair process^38, 39^.

To address this, we used an *in vivo* model of bone injury to longitudinally measure changes in pro-and anti-inflammatory monocytes and macrophages in addition to osteoblast and osteoclast numbers and bone volume at the site of injury. We then used the obtained biological data combined with empirically-derived parameters from the literature to power an ODE model of bone injury repair and examine the impact of infiltrating immune cells on osteoblast and osteoclast activity over time in regard to bone volume dynamics. The ODE model generated herein, demonstrated that the temporal interplay between myeloid derived pro- and anti-inflammatory populations are critical in driving osteoblast and osteoclast response but interestingly, using a constant rates of bone formation and resorption, the mathematical model failed to recapitulate the bone volume dynamics. Further interrogation of the model demonstrated that the rate of osteoclast resorptive activity must vary greatly over the course of injury resolution to return the bone volume to homeostasis. This insight has not been considered to date and underscores the value of mathematically modeling complex multicellular biological process.

## RESULTS

### Osteoclast and osteoblast numbers fluctuate dynamically in response to bone injury

The stages and duration of non-critical bone injury largely follow the same program, whereby subsequent to injury, early inflammation and hematoma occur rapidly, followed by the formation of a callus that is subsequently mineralized by bone-forming osteoablasts^4, 5, 40^. The callus is then remodeled via the activity of bone resorbing osteoclasts^1, 18, 41–46^ (Fig. 1a). Osteoblasts and osteoclasts are critical mediators of these steps and their numbers shift accordingly during each phase of repair. Existing theoretical models of bone remodeling assume osteoblast and osteoclast activities are constant over time and therefore, their numbers directly predict bone dynamics^31, 32, 34, 37^. To evaluate this prevailing assumption, we first asked if modeling osteoclast and osteoblasts alone was sufficient to accurately predict corresponding bone remodeling dynamics using experimental data. To generate parameters to power such an ODE model, we used an experimental model of bone injury repair: non-critical injury resulting from direct intratibial penetration via the knee epiphysis into the medullary canal^47–50^ (Fig. 1b). Tibias from mice were collected prior to injury at baseline (day 0), and at day 1, 2, 3, 7 and 14 (n = 5 mice/time point) following injury. High-resolution μCT analysis of uninjured tibia established baseline bone volume (BV/TV) (Fig. 1c and d). Our data show that after injury, bone volume around the injury site diminished over a 48-hour period, prior to a robust increase in mineralized bone content between days 2 and 7. By day 14, the bone volume returned toward baseline values. We directed our μCT and histological analyses on the area surrounding the bone injury rather than the entire bone marrow since our goal was to quantify cellular dynamics and changes specifically in response to injury; values that could be diluted by measurements in non-injured areas of the medullary canal (Supplemental Fig. 1). Focusing on the site of injury and surrounding area, histologically, we observed sequential increases in osteoblasts followed by osteoclasts, findings that are qualitatively consistent with our BV:TV μCT analyses and are in line with previous published observations^50–52^ (Fig. 1c and d).

**Fig. 1.**
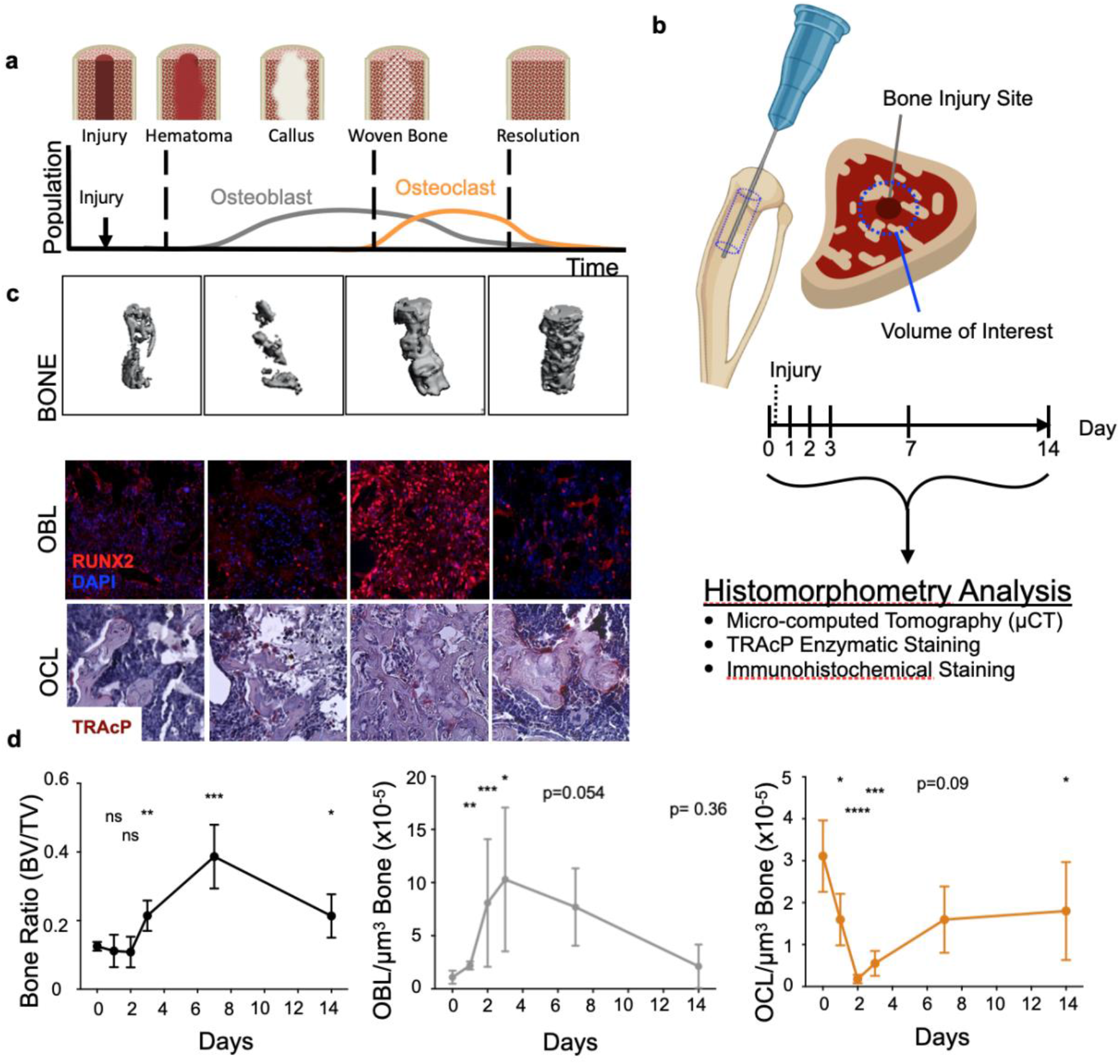
Osteoblast (OBL) and osteoclast (OCL) numbers temporally fluctuate dynamically as bone heals from injury. **a** schematic summarizing published dynamics of OBL and OCL following bone injury. **b** schematic depicting the experimental workflow to induce bone injury in mice and generate bone, osteoblast and osteoclast dynamic data. **c** representative images of micro-computed tomography revealed trabecular bone status (BONE). Decalcified bones were stained and quantified for OBL by RUNX2 immunofluorescence staining (OBL), and OCL by tartrate-resistant acid phosphatase (TRAcP) staining (OCL). **d** quantitation of temporal dynamics of bone volume, osteoclast and osteoblast population.

### Bone repair dynamics cannot be computationally recapitulated using constant osteoblast and osteoclast activity rates

To date, bone resorption and formation rates have been difficult to measure *in vivo*. Despite various *in vitro* studies showing that osteoblast and osteoclast activity can be controlled by inflammatory factors and cytokines^15, 53–64^, existing theoretical mathematical models of bone remodeling largely assume that resorption and formation rates per cell are fixed/constant over time. Since measuring whether osteoblast and osteoclast activities vary over time *in vivo* during bone injury repair is experimentally challenging, we employed an integrated experimental and mathematical approach to address this knowledge gap.

Using the obtained biological data and publicly-available parameter values regarding osteoblast and osteoclast behavior (Fig. 1c, Supplemental Fig. 2 and Table 1), we developed an initial mathematical data-driven ODE model to recapitulate the control of bone repair exclusively by these two populations (Fig. 2a). This initial ODE model simulated the bone injury event as a transient osteoblast (OBL) expansion and a decrease in osteoclast (OCL) population from day 0 to 2 (see mathematical and computational methods). Fits to the rest of the OCL and OBL population data were optimized within the parameter space defined by published literature, such as regarding cellular lifespan and proliferation rates (Fig. 2b and Table 1). Optimal fits with greatest R^2^ value and number of residuals less than 1 (#R<1) were subsequently used for estimating bone volume dynamics. Using these OCL and OBL optimized fits, the ODE model attempted to recapitulate experimental bone dynamics by sampling *constant* bone resorption rates within a range previously described in literature^50, 53, 65, 66^. A corresponding bone formation rate was estimated in each sampling as to ensure a return to baseline bone volume at the end of the injury repair process (Fig. 2a **#**). Interestingly, using this iterative approach, the ODE predictions largely overestimated the bone volume dynamics compared to the experimental data (Supplemental Fig. 3). In fact, the best-fitted iteration, that used the lowest published OCL resorption rates^66^, only achieved an R^2^ value of 0.4554, and #R<1 of 2/5 (Fig. 2c). This indicated that either published measurements of *in vitro* bone resorption/formation parameters do not reflect *in vivo* rates, and/or that bone resorption/formation rates by osteoclasts and osteoblasts are variable over time during the course of injury repair. To address this, we alternatively fitted the model to bone dynamics data while allowing the optimization algorithm to freely determine an optimal combination of *constant* bone resorption and formation rates that were not forced to return to baseline bone volume subsequent to injury (Fig. 2a **&**). This resulted in improved bone volume dynamic fits during injury repair but, of note, the final bone volume reached by the ODE was 70% lower compared to that of baseline (Fig. 2d). Taken together, these data suggest that osteoclast and osteoblast activity rates must vary greatly during injury response in order to return the bone to homeostasis during injury repair time-frame. This raised the question as to what cellular/environmental cues are potentially responsible for controlling their activity.

**Fig. 2.**
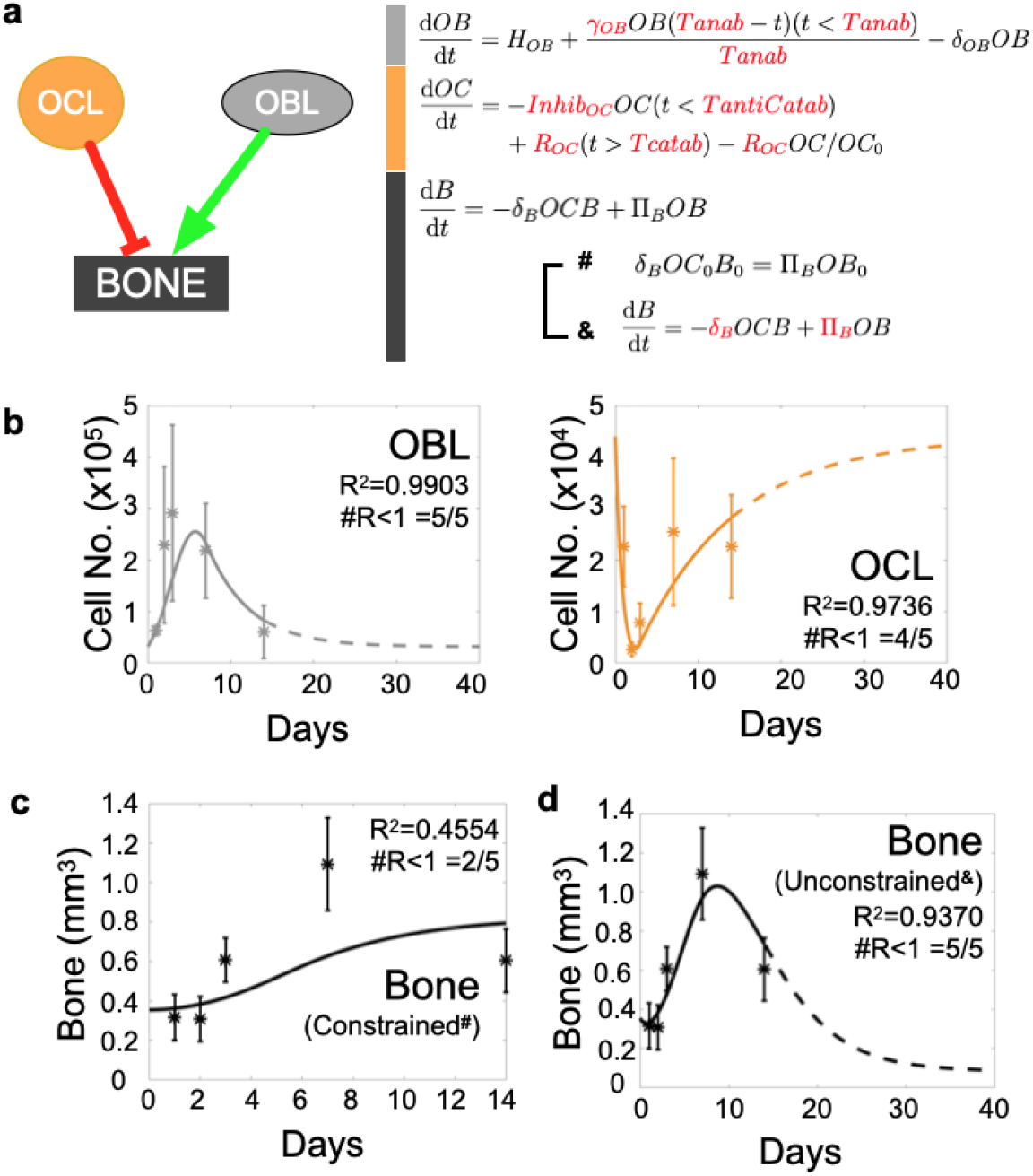
Osteoblast (OBL) and osteoclast (OCL) activities as measured at homeostasis do not allow accurate bone prediction during injury repair *in vivo*. Histological quantitation of tibia bones parameterizes mathematical ordinary differential equation (ODE) model of bone injury repair. **a** ordinary differential equations (ODE) describing dynamics of OCL and OBL population are paired with published parameters to form an initial ODE model to predict bone repair dynamic. Schematic depicts OCL resorb (red line) and OBL form bone (green line). ODE expressions with unknown value (red) were estimated as the model optimizes fits to *in vivo* data. **b** model produces accurate fits to OCL and OBL dynamics. **c** model falsely predicts bone dynamics given OBL and OCL fits in the first 14 days following bone injury when it samples various publication-derived OCL resorption rates (each dashed line represents one sampling). OBL bone formation rates are mathematically estimated in each sampling to ensure predictions will eventually return to homeostasis (**#**). **d** alternatively, ODE model was allowed to freely seek out a combination of resorption and formation rates to best fit data within the 14-day time period (**&**).

**Table 1.** Parameters extracted from published literature were combined with temporal dynamics data in ODE model to estimate previously unknown parameters needed to fit bone data. apposition rates from confined model (**#**) were calculated in fashion to offset resorption rates derived from publication, to maintain constant bone volume at homeostasis.

| Parameter | Description | Value | Unit | Supplemental References | Standard Error |
| --- | --- | --- | --- | --- | --- |
| $\delta_{OB}$ | Osteoblast Lifespan | 0.19 | Day<sup>-1</sup> | 33, 34 | -- |
| $\gamma_{OB}$ | Osteoblast Proliferation Rate | 0.873 | Day<sup>-1</sup> | Estimated | 0.1999 |
| $Inhib_{OC}$ | Osteoclast Inhibition | 1.2186 | Day<sup>-1</sup> | Estimated | 0.3540 |
| $R_{OC}$ | Osteoclast Recruitment | 6774.8 | Cell Day<sup>-1</sup> | Estimated | 1967.6 |
| $T_{anab}$ | Duration of Anabolism | 6.6924 | Day | Estimated | 1.8649 |
| $T_{antiCatab}$ | Osteoclast Inhibition Duration | 2 | Day | Estimated | -- |
| $T_{Catab}$ | Starting Time of Catabolism | 2 | Day | Imposed | -- |
|  | Published Resorption Rate Range<sup>#</sup> | [1 X 10<sup>-8</sup>-5 X 10<sup>-5</sup>] | mm<sup>3</sup> Cell<sup>-1</sup> Day<sup>-1</sup> | 26, 27, Imposed | -- |
| $\delta_B$ | Per Bone Volume Unit Resorption Rate<sup>#</sup> | 7.034 X 10<sup>-6</sup> | Cell<sup>-1</sup> Day<sup>-1</sup> | Estimated | 7.30 X 10<sup>-7</sup> |
| $\Pi_B$ | Bone Formation Rate<sup>#</sup> | -- | mm<sup>3</sup> Cell<sup>-1</sup> Day<sup>-1</sup> | Determined from $\delta_B$ | -- |
| $\delta_B$ | Per Bone Volume Unit Resorption Rate<sup>&</sup> | 5.7687 X 10<sup>-6</sup> | Cell<sup>-1</sup> Day<sup>-1</sup> | Estimated | 1.63 X 10<sup>-6</sup> |
| $\Pi_B$ | Bone Formation Rate<sup>&</sup> | 1.2164 X 10<sup>-6</sup> | mm<sup>3</sup> Cell<sup>-1</sup> Day<sup>-1</sup> | Estimated | 1.6043 X 10<sup>-7</sup> |

### Polarized pro- and anti-inflammatory monocytes and macrophages emerge in distinct temporal waves during bone injury repair

Monocytes and macrophages are key cellular species in the bone ecosystem and their pro- and anti-inflammatory functions have been implicated in the bone injury repair process and in the regulation of osteoblast/osteoclast activity^8, 18, 19, 50, 67, 68^. Studies have shown for example that, 1) myeloid cells are polarized in bone injury and inflammation, 2) pro-inflammatory factors and myeloid cells stimulate osteoclast activity, and 3) anti-inflammatory/wound-healing factors and myeloid cells stimulate osteoblast activity (Supplemental Fig. 4)^1, 9, 11, 45, 51, 67–69^. Based on this rationale, we therefore hypothesized that fluctuations in the number and polarization status of myeloid populations control osteoclast and osteoblast activity during bone repair. To test this hypothesis, we reanalyzed the non-critical bone injury experiment. Tibias from mice were collected at baseline prior to injury (day 0), and at day 1, 2, 3, 7 and 14 (n = 5/time point) post-injury. Flow cytometry was used to measure changes in myeloid populations over time^8, 17, 70–80^ (Fig. 3a-c and Supplemental Fig. 5). Our results show that there are significant increases in pro-inflammatory monocytes and macrophages within the first 48 hours that are subsequently rapidly depleted upon the infiltration of anti-inflammatory macrophages between 24 and 72 hours (Fig. 3c). Interestingly, in accordance with observations from other *in vivo* studies, we observed a smaller second wave of pro-inflammatory monocytes between days 6 and 8^81–83^ (Fig. 3c and d).

**Fig. 3.**
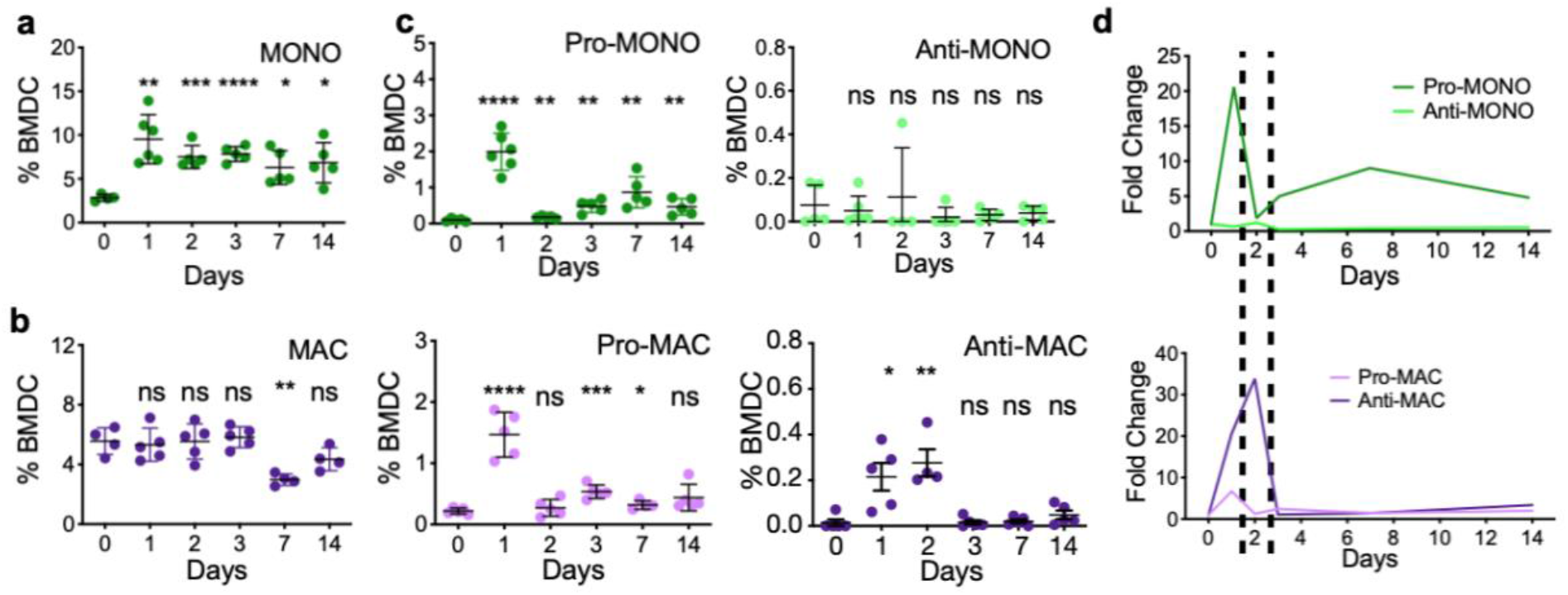
Transient waves of pro- and anti-inflammatory monocytes and macrophages alternate dynamically during bone injury repair. Flow cytometry performed on tibia bone marrow harvested from C57BL/6 mice at various time points after injury (n=30; 5/ time point) reveals diverse myeloid dynamics and polarization. Time points corresponds to time points from histological data. Total monocyte (CD11b+ LY-6C^HI^ LY-6G-; **a**) and macrophage (CD11b+ LY-6C^LO^ LY-6G-; **b**) and their respective pro- and anti-inflammatory subsets (**c**) each uniquely fluctuates following bone injury (Student t-test compares all time points to its Day 0 for each subset; *p<0.05 **p<0.005 ***p<0.0005 ****p<0.00005 ^ns^p>0.05). **d** temporal dynamics of pro and anti-inflammatory monocytes and macrophage numbers are normalized as fold change relative to levels at homeostasis. Dashed lines show timings of pro- and anti-inflammatory polarization are mutually exclusive.

### Integration of polarized myeloid cells control of bone remodeling activity recapitulates bone healing dynamics

Previous studies have reported pro- and anti-inflammatory myeloid control of osteoclast and osteoblast activity; however, these observations are largely derived from *in vitro* settings^54, 67, 84–93^. To address this, we integrated the experimental quantitative data collected from each of these populations via flow cytometry into the framework of the ODE model (Fig. 4a and b). Specifically, we allowed osteoclast activity to be stimulated from baseline in proportion to the presence of pro-inflammatory cells by a model-estimated constant factor of α. Likewise, we allowed osteoblast activity to be stimulated from baseline in proportion to the presence of anti-inflammatory cells by a model-estimated constant factor of β. These assumptions are based on empirical data from published *in vitro* experimental data^54, 67, 84–86, 88–93^. We then asked the expanded ODE model to optimize for levels of α and β that are needed to recapitulate bone volume dynamics. Of note, we did not integrate anti-inflammatory monocyte data as the experimental data demonstrated this population remains consistently low levels that did not fluctuate throughout the course of bone injury repair (Fig. 3c). Importantly, in our model optimization, the range of osteoblast and osteoclast activities that could be influenced by infiltrating myeloid cells were limited to published values (Supplemental Fig. 2). Given these restraints, the model nevertheless estimated an optimal set of parameters that significantly recapitulated the bone volume dynamics (R^2^= 0.9362; #R<1 =5/5) (Fig. 4c). The optimized model reveals that while osteoblast activity remains relatively constant, osteoclast activity changes dramatically over time. Furthermore, in this expanded ODE model, the bone volume returned to baseline levels subsequent to injury, underscoring the biological validity of our model assumptions and reinforcing the importance of myeloid-derived infiltrating cells in controlling the activity of osteoblasts and in particular osteoclasts in the process.

**Fig. 4.**
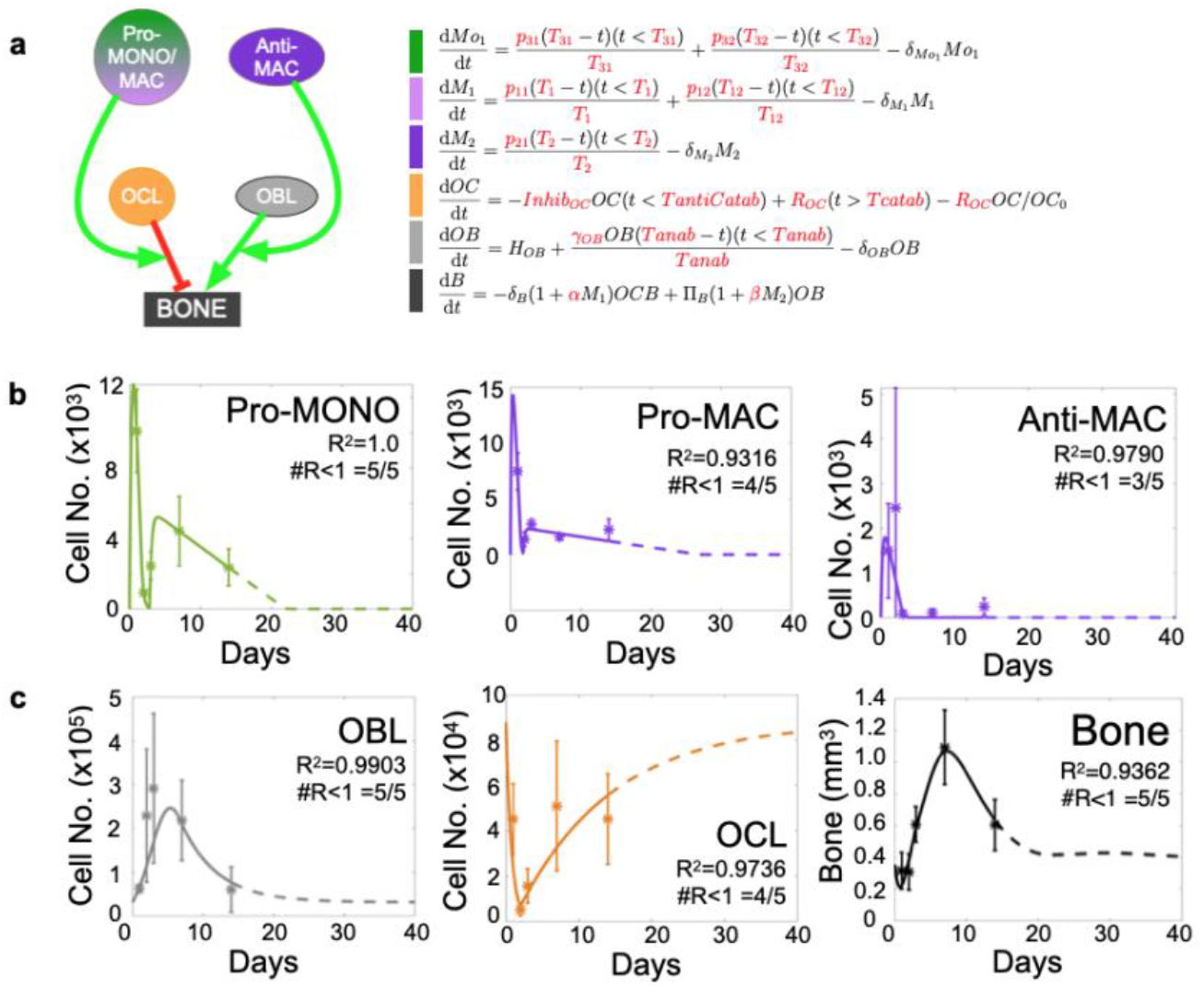
Integration of pro- and anti-inflammatory myeloid populations to modulate OCL and OBL activity sufficiently improves model fit to experimental bone data. **a** ODE expands to six populations and allows manually-fitted pro- and anti-inflammatory cells to enhance bone resorption and formation rates, respectively. Individual equations are shown next to schematic of ODE framework, model estimates amount of influence polarized myeloid cells have on bone remodeling activity to optimize fit to data (expressions in red). **b** Manual fits to pro- and anti-inflammatory monocytes (Pro- and Anti-MONO, respectively), anti-inflammatory macrophages (Anti-MAC) are represented by solid lines through error bar of data. **c** myeloid data was used to predict bone dynamics given OBL and OCL fits. Statistical analysis of resulting fits on OCL, OBL and bone are shown (R^2^).

### Osteoclast resorption activity does not correlate with osteoclast number during bone injury repair

Upon further analysis of the results generated by our expanded ODE model, we noted a disconnect between the dynamics of osteoblast and osteoclast activity versus their population numbers (Fig. 5a). The model predicts that osteoblast mineralizing activity varies slightly over time 1.21×10^-6^ to 2.63×10^-6^ mm^3^/cell/day; however, the model predicts that a range of 4.26×10^-7^ to 7.28×10^-6^ mm^3^/cell/day is required for osteoclast activity (Fig. 5a and b). These data suggest that, while osteoblast activity only increases by 2 folds, a 17-fold increase in osteoclast activity is required to recapitulate injury dynamics and also return to the bone volume to homeostasis. Importantly, the noted ranges for osteoclast activity fall within those values reported in independent studies^53, 65, 66^ (Fig. 5b and Supplemental Fig. 2). Our model is the first to posit that the rate at which osteoclast resorbs mineralized matrix in bone healing can vary greatly depending on cues from the surrounding microenvironment.

**Fig. 5.**
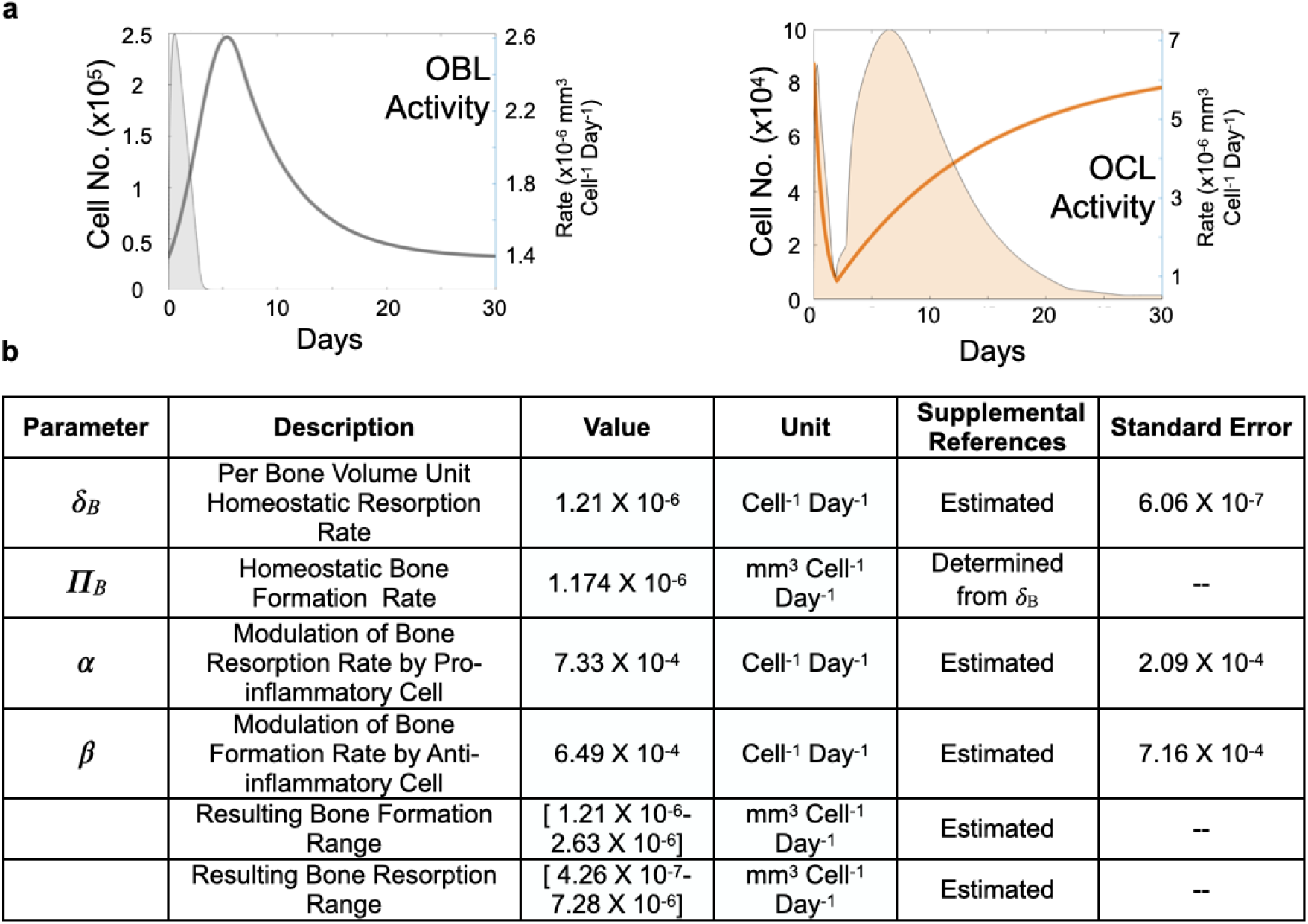
OCL and OBL activities and numbers do not correlate and vary distinctly across in bone injury repair. **a** OBL and OCL activity rate dynamics (filled curves plotted on the right y-axis) are plotted against their population dynamics (unfilled curves plotted on the left y-axis). Activity is temporally distinct from population dynamics, both combine to recapitulate bone dynamics. **b** table detailing the known parameters used by model to estimate/infer unknown parameters.

We also submit that this variation in resorptive activity is driven by infiltrating pro- and anti-inflammatory monocytes and macrophages. To this end, we returned to our initial ODE model (Fig. 2) and asked how would osteoclast activity change overtime in order to fit to the bone data. In this agnostic approach, we no longer restricted osteoclast resorption to constant rates over time, but defined a piecewise linear function of time for osteoclast resorption rate (Supplemental Fig. 6a and b, Mathematical and Computational Methods). Independent of the parameters chosen for the initial piecewise slope conditions, the optimization algorithm identified a functional form composed of two waves: initially intense and transient between days 1 and 2, followed by a milder but persistent wave starting after Day 3 (Supplemental Fig. 6c; orange line). We noted this temporal profile was very similar to that of experimental data regarding the pro-inflammatory monocytes and macrophages populations (Fig. 3c and Supplemental Fig. 6c). This result further supports that pro-inflammatory cells contribute significantly to osteoclast behavior and therefore bone healing dynamics (Supplemental Fig. 6). We used a similar approach to determine variable osteoblast activity and defined a piecewise linear function of time, for the bone formation rate (Supplemental Fig. 7a). Interestingly, this model did not recapitulate bone volume dynamics despite the freedom to change bone formation rate over time (Supplemental Fig. 7b; R^2^=0.6796). These data further support a role for pro-inflammatory myeloid cells in controlling osteoclast, and less so osteoblast activity, and therefore bone volume during injury repair.

## DISCUSSION

The complex cellular mechanisms that control bone injury repair can be difficult to dissect given the complexity of the bone marrow microenvironment using traditional biological approaches but key insights have been made. For example, genetic and pharmacologic approaches reveal that macrophages play important roles in bone healing as well as osteoclast differentiation^51, 68, 93–95^. Yet, how macrophage populations quantitatively interact with each other or other cell types in the bone environment directly or indirectly over time can be challenging to identify with this approach. For instance, though polarized macrophages have been observed at sites of bone injury alongside osteoblasts and osteoclasts, the rates at which polarized macrophages stimulate the activities of these bone cells remained difficult to evaluate and furthermore, quantitate. Computational approaches allow simultaneous interrogation of multicellular systems in which a mathematical model can infer parameter values that may be otherwise unknown. Despite this advantage, existing mathematical models of bone remodeling largely focus only on osteoclast, osteoblast and the bone; and those which integrate additional populations are theoretical^28, 30–38^. As shown here, we have integrated both our experimental and published data into a mathematical framework which models interactions between myeloid cells, osteoclasts and osteoblasts, and the bone from published literature. We have concluded that bone repair cannot be recapitulated if we assume osteoclast osteoblast activities are constant over time. Our initial ODE model failed to derive accurate bone fits despite using both published and freely estimating *constant* activity rates. Therefore, based on the literature, we subsequently focused on myeloid-derived monocytes and macrophages that have noted roles in contributing to bone injury repair^51, 68, 93–95^, and the expanded ODE demonstrated that the dynamic waves of polarized monocytes and macrophages and their temporal control of bone remodeling activity sufficiently allowed for the accurate recapitulation of bone repair.

Another major finding from our analyses is the extent to which osteoclast resorptive activity can be modulated subsequent to osteoblast mineralization of the injury site. Existing empirical data have recorded osteoclast resorptive activities in the range of 1×10^-8^ to 5×10^-5^ mm^3^/cell/day^53, 65, 66^. Here, our estimations suggest that osteoclast activity varies by 17-fold magnitude over time within this published range and that pro-inflammatory monocyte and macrophages are critical for regulating this effect. These data underscore how mathematical modeling can provide important biological insights. It should be noted that though other mathematical models have been proposed to explore mechanisms of bone repair dynamics^30, 32, 34, 36, 38^, the study presented herein, to our knowledge, is the first to leverage longitudinal biological data on multiple cellular populations and integrate this information into a mathematical model.

Additional quantitative insights provided by the ODE model include estimations on monocyte and macrophage proliferation rates as well as the rates at which pro- and anti-inflammatory cells polarize and modulate osteoclast and osteoblast activity, respectively, during the repair process. This information can be critical for therapies that target specific myeloid populations during bone injury repair in a bid to accelerate bone healing. Our study also reveals rapid expansion of pro-inflammatory monocytes and macrophages in the first 24 hours with anti-inflammatory macrophages emerging shortly thereafter and persisting for up to 48 hours. Interestingly, pro-inflammatory cells moderately rebound upon the clearance of anti-inflammatory cells (between days 6 and 8, Fig. 3c), suggesting a second wave of inflammation that is in keeping with other reports^81–83^. Conflicting reports suggest this could be due to 1) emergence of anti-inflammatory macrophages having an inhibitory effect on pro-inflammatory population, or 2) myeloid plasticity and repolarization^68, 72, 96–101^. Our next efforts with the ODE generated herein will focus on the interplay between macrophages and how their polarization states control not only each other, but also how osteoblasts and osteoclasts coordinate bone injury repair.

One caveat of our study is that the flow cytometric analysis is performed on cells isolated from the whole bone marrow, as opposed to only the volume of interest in histological datasets (See Supplemental Fig. 1). An alternative could be to perform multiplex image cytometry of the site of injury for the various myeloid populations of interest. We suspect that, while this would allow for more accurate quantitation of the myeloid cell populations infiltrating the site of information, the overall trends and shifts in those populations over time would remain similar to our flow cytometry data. Additionally, our model does not consider the potential roles of other cell types in the bone ecology that could contribute, such as T cells. Our results suggest that modeling myeloid populations provides enough resolution to satisfactorily explain the process of non-critical bone injury repair. Importantly, our unbiased test (Supplemental Fig. 6 and 7) yielded osteoclast activity dynamics that qualitatively resonated with the population dynamics of pro-inflammatory monocytes and macrophages, supporting the importance of the myeloid population in regulating bone volume resorption. Our theoretical framework is flexible enough however that the effects of other immune cells such as T cells could be included in future iterations of our ODE model.

Through our unbiased data-driven testing approach, we have integrated experimental data into a physiologically-relevant mathematical model exploring non-critical bone injury repair. A potential application of our modeling approach is to determine how bone healing times subsequent to injury can be improved via therapeutic intervention. Previous reports have demonstrated that modulating pro- and anti-inflammatory macrophages can alter the time taken for bone injury repair^14, 19, 50, 89^. Because of the ability of the ODE model to recapitulate the temporal dynamics of the cellular populations involved in bone injury repair, we can investigate the precise timing at which to administer therapies in order to further shorten bone-healing time. Likewise, we can examine cellular behavior in response to a different sized injury, or even in a different bone injury context, such as non-union fractures. These points will be best addressed once we are able to enhance our ODE model with reciprocal mechanisms and fully couple the system.

In conclusion, we have developed an ordinary differential equation (ODE) model of osteoclast, osteoblast and bone dynamics, that considers the influences of polarized myeloid cells during bone injury. The model faithfully recapitulates bone volume dynamics during injury repair and returns to homeostasis. It further yields a number of novel insights regarding myeloid control of osteoclast- and osteoblast-mediated bone resorption and formation over time. To our knowledge, this model is the first to recapitulate longitudinal in vivo data of simultaneously measured bone and myeloid cell populations, as well as bone volume during bone healing. A better understanding of bone healing will have clinical translatability, allowing, for instance, accelerating the process and improve patient outcomes.

## MATERIALS AND METHODS

### Intratibial Bone Injury Model

All animal studies were performed in accordance with Guidelines for the Care and Use of Laboratory Animals published by the National Institutes of Health, under IACUC Protocol R5857 (CCL). Additionally, studies abided by relevant ARRIVE guidelines. 5-6-week-old male immune-competent C57BL/6 mice were purchased from Jackson Laboratory with consideration for study statistical significance and power (n=30). Surgically prepared mice (n=25) were sterilized with chlorhexidine and subject to non-critical bone injury by intratibial injection using a 28-gauge (0.3062mm diameter) syringe by penetration through the knee epiphysis to mid-shaft. Five mice remained uninjured and were euthanized at baseline, and, subsequently, randomly selected injured mice were euthanized at days 1, 2, 3, 7 and 14 (n=5/time point) for histological and flow cytometry analyses. Histological and FACS data were obtained in a blind manner to parameterize subsequent mathematical models.

### Micro-Computed Tomography

Injured tibias harvested from mice from all time points were centralized and were subjected to micro-computed topography (μCT) scanning using Scanco μ35 scanner to derive bone volume data. Individual bone scans were deidentified using numerical codes during, and reidentified following data analysis in a blinded fashion. A gap of 100μm from the tip of growth plate towards the midshaft was avoided to ensure the high bone density nature of the growth plate does not mask potential differences in bone volume associated with the injury. Each bone was then scanned every 6μm for a total span of 1000*μ*m along the midshaft. Trabecular bone histomorphometry was subsequently performed after contouring each slice scan and reconstructing the 3-dimension volume of interest structure of each bone using the built-in morph function (n=30 bones; 5/time point). This process was performed repeatedly using different contours to generate bone status dynamics of the whole trabeculae, the region surrounding the injury, and of the injury itself (Supplemental Fig. 1).

### TRAcP Staining

Tibia bones from all time points were decalcified with 14% EDTA every other day for 3 weeks for further staining quantitation and analyses following μCT scans. Formalin fixed paraffin embedded (FFPE) bones were sectioned at 4μm thickness. Multiple slides sectioned at different depths from each bone were pooled for all time points, and were baked at 42°C overnight to improve adhesion while retaining enzymatic activity for tartrate-resistant acid phosphatase (TRAcP) enzyme-based staining for osteoclast numbers. Deparaffined and rehydrated sections were pre-incubated in basic stock solution with napthol-ether substrate for 1 hour at 37°C and developed in pararosaniline dye and sodium nitrite for 10mins, also at 37 °C. Sections with red osteoclasts were further counterstained with hematoxylin to visualize bone tissue morphology. Fixed slides were imaged at 20X using Evos Auto brightfield microscopy to include injury site and its immediate periphery. All TRAcP positive (red) multinucleated osteoclasts within 5*μ*m radius from injury were counted, and mathematically converted to osteoclasts / bone marrow volume (#OCL/μm^3^) for each slide for each bone at each time point. This region of is consistent in area with the μCT analysis parameters to ensure consistency in data.

### Immunofluorescence Staining and Quantitation

Additional FPPE tibia bone sections were baked at 56°C in preparation for immunofluorescence staining of osteoblast (RUNX2 at 1:500; Abcam Cat. No. ab81357) and nuclear staining (DAPI). Slides were processed in batch similar to TRAcP staining methodology. Deparaffined and rehydrated slides were subject to heat-induced antigen retrieval method. Sections were then blocked and incubated in primary antibodies diluted in 10% normal goat serum in TBS overnight at 4°C. Subsequently, slides were stained with secondary Alexa Fluor 568-conjugated antibody at 1:1000 at room temperature for 1 hour under light-proof conditions. Stained slides were stained with DAPI for nuclear contrast and mounted for imaging at 20X using Zeiss upright fluorescent microscope to include the injury site as well as the immediate peripheral tissue. All RUNX2 positive cells (red staining colocalizing with DAPI) within 5*μ*m radius from injury were counted and mathematically converted to osteoblasts / bone marrow volume (#OBL/μm^3^) for each bone at each time point. Again, this methodology ensured consistency across all acquired datasets.

### Flow Cytometry and Analysis

Harvested contralateral injured tibias (n=30; 5/time point) had ends removed and were subjected to centrifugation at 16,000g for 5 seconds for isolation of whole bone marrow for flow cytometry staining and analysis. Red blood cells were lysed using RBC Lysis Buffer from Sigma Aldrich (Cat. No. R7757-100ML) as per manufacturer’s guidelines. Live bone marrow cells were subject to FcR-receptor blocking (1:3; BioLegend; Cat. No. 101319) and viability staining (1:500; BioLegend; Cat. No. 423105). Samples were then stained by cell-surface conjugated antibodies from BioLegend diluted in autoMACS buffer (Miltenyi; Cat. No. 130-091-221) for phenotyping myeloid cells: CD11b-BV786 (1:200; Cat. No. 101243), LY-6C-Alexa Fluor 488 (1:500; Cat. No.128021) and LY-6G-Alexa Fluor 700 (1:200; Cat. No. 561236). Cells were then fixed with 2% paraformaldehyde in dark prior to intracellular staining. Fixed cells were permeabilized using intracellular conjugated antibodies to assess polarization status: NOS2-APC (1:100; eBioscience; Cat. No. 17-5920-80) and ARG1-PE (1:100; R&D; Cat. No. IC5868P). Appropriate compensation and fluorescence-minus-one (FMO) controls were generated in parallel either with aliquots of bone marrow cells or Rainbow Fluorescent Particle beads (BD Biosciences; Cat. No. 556291). All antibody concentrations were titrated prior to injury study using primary bone marrow cells to ensure optimal separation and detection of true negative and positive populations. Stained controls and samples were analyzed using BD Biosciences LSR flow cytometer (Supplemental Figure 4). All datasets were batch analyzed to ensure optimal consistent gating stringency.

## MATHEMATICAL AND COMPUTATIONAL METHODS

### Model Parameterization

#### Initial ODE model

The initial Osteoblast/Osteoclast/Bone ODE model is presented in equation Fig. 2a. OB, OC and B represent osteoblasts, osteoclasts and bone volume, respectively. Osteoblast and osteoclast equations are composed of a homeostatic source term, a clearance term, and an injury-triggered expansion term. The osteoblast clearance parameter δ_OB_ was fixed from literature and in order to ensure osteoblast homeostasis level, the source term H_OB_ was fixed at δ_OB_*OB_0_, where OB_0_ represents the initial level of osteoblasts. The osteoblast proliferation rate γ_OB_ and duration of expansion T_anab_ were calibrated in fitting the osteoblast dynamics to the experimental data. The osteoclast decrease rate (Inhib_OC_), the decrease duration (T_antiCatab_), the replenishment time (T_Catab_) and the replenishment rate (R_OC_) were calibrated in fitting the osteoclast dynamics to the experimental data. The homeostatic clearance parameter was fixed to R_OC_/OC_0_ in order to ensure homeostasis back to the initial osteoclast level.

The bone equation comprises two terms: a bone resorption term, proportional to the number of osteoclasts and proportional to bone volume, and a bone formation term, proportional to osteoblast number. The resorption term is proportional to bone volume since less bone translates to less bone available for osteoclast resorption. On the other hand, osteoblast-mediated bone formation is independent of available bone. A range of possible resorption rates was derived from published measurements. Equation on Fig. 2a **#** shows how bone formation parameter is fixed at δ_B_OC_0_B_0_/OB_0_, where B_0_ is the initial bone level, in order to ensure that bone level remains at homeostasis when osteoclast and osteoblast levels are at homeostasis (Corresponding predictions on Fig. 2c). Equation on Fig. 2a **&** shows the case where both δ_B_ and Π_B_ are freely optimized (corresponding fit on Fig. 2d).

#### Piecewise linear temporal variation of bone resorption and formation rates

The initial ODE model was further used to study time-dependent bone resorption rate. It was defined as an explicit piecewise linear function of time (Supplemental Fig. 6a and 7a). The dynamics of osteoclast and osteoblast was imposed from the previous fit. The successive slopes of the piecewise linear function and the initial resorption rate were all estimated from fitting the resulting bone dynamics to bone experimental data. The parameter space for the optimization was defined as follows: The slopes were allowed to be positive or negative, with the constrain that the resorption rate cannot become negative or go beyond the upper bound defined from literature. Using the same approach, model sought to recapitulate bone dynamics by optimizing osteoblast activity dynamics (Supplemental Fig. 7).

#### Enhanced ODE model including polarized monocytes/macrophages

For the polarized macrophage part (Fig. 4), cells clearances/lifespans were fixed from literature and all the other parameters were calibrated to fit the experimental data. The osteoblast and osteoclast fits were kept the same as in the initial model. For the bone equation, homeostatic bone resorption rate δ_B_ (before and after injury), pro-inflammatory monocytes/macrophages-mediated bone resorption stimulation parameter (α) and anti-inflammatory macrophages-mediated bone formation stimulation parameter (β) were all calibrated to fit the experimental bone dynamics. The homeostatic bone formation rate Π_B_ was fixed such that Π_B_ =δ_B_OC_0_B_0_/OB_0_ so bone level is ensured to remain at homeostasis when osteoclast and osteoblast levels are at homeostasis in absence of injury (no polarized monocytes and macrophages).

### ODE Solver

The ODE45 function of Matlab was used to solve the differential equation system. The experimental baseline values (time 0) were used as initial conditions.

#### Parameter estimation method

To estimate parameters facilitating goodness of fit, we defined the following objective function:

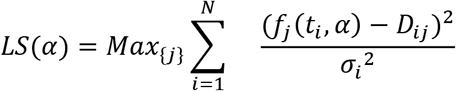

Where i represents the time point index and j the variable index, *α* represents the parameter set used to evaluate the model function f, *D_ij_* represents the experimental data of variable j at time point i, and *σ_i_* represents the experimental error. The choice of this functional form instead of the sum of the squares of the residuals was motivated to avoid that one fit variable would be “sacrificed” to the benefit of another one. This way, we ensure that all variables are equally well fitted.

- In order to minimize this function representing the error estimate between data and model, we used the Matlab function fminsearch with a penalization term to stay in a parameter range set with reasonable boundaries.
- AIC criterion is defined as follows:

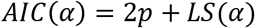 Where p is the number of parameters.

## AUTHOR’S CONTRIBUTIONS

C. H. Lo, E. Baratchart. C. C. Lynch and D. Basanta were responsible for study conception and design. C. H. Lo, E. Baratchart. C. C. Lynch and D. Basanta developed the methodology. C. H. Lo and E. Baratchart. acquired the data while C. H. Lo, E. Baratchart. C. C. Lynch and D. Basanta were responsible for analysis and interpretation of data. C. H. Lo, E. Baratchart. C. C. Lynch and D. Basanta were responsible for manuscript writing and editing.

## DISCLOSURES OF CONFLICT OF INTEREST

The authors have no potential conflicts of interest to disclose.

## Notes

### Competing Interest Statement

The authors have declared no competing interest.

## REFERENCES

1. Schindeler, A., McDonald, M.M., Bokko, P. & Little, D.G. Bone remodeling during fracture repair: The cellular picture. Seminars in cell & developmental biology 19, 459–466 (2008).

2. Gerstenfeld, L.C., Cullinane, D.M., Barnes, G.L., Graves, D.T. & Einhorn, T.A. Fracture healing as a post-natal developmental process: molecular, spatial, and temporal aspects of its regulation. J Cell Biochem 88, 873–884 (2003).

3. Gwathmey, F.W., Jr., Jones-Quaidoo, S.M., Kahler, D., Hurwitz, S. & Cui, Q. Distal femoral fractures: current concepts. J Am Acad Orthop Surg 18, 597–607 (2010).

4. Chen, W.T. et al. A special healing pattern in stable metaphyseal fractures. Acta Orthop 86, 238–242 (2015).

5. Uhthoff, H.K. & Rahn, B.A. Healing patterns of metaphyseal fractures. Clin Orthop Relat Res, 295–303 (1981).

6. Atanelov, Z. & Bentley, T.P. Greenstick Fracture, in StatPearls (StatPearls Publishing Copyright © 2020, StatPearls Publishing LLC., Treasure Island (FL); 2020).

7. Bahney, C.S., Hu, D.P., Miclau, T., 3rd & Marcucio, R.S. The multifaceted role of the vasculature in endochondral fracture repair. Frontiers in endocrinology 6, 4 (2015).

8. Horwood, N.J. Macrophage Polarization and Bone Formation: A review. Clinical reviews in allergy & immunology 51, 79–86 (2016).

9. Loi, F. et al. Inflammation, fracture and bone repair. Bone 86, 119–130 (2016).

10. Cho, S.W. et al. Osteal macrophages support physiologic skeletal remodeling and anabolic actions of parathyroid hormone in bone. Proceedings of the National Academy of Sciences of the United States of America 111, 1545–1550 (2014).

11. Wu, A.C., Raggatt, L.J., Alexander, K.A. & Pettit, A.R. Unraveling macrophage contributions to bone repair. BoneKEy reports 2, 373 (2013).

12. Chan, J.K. et al. Low-dose TNF augments fracture healing in normal and osteoporotic bone by up-regulating the innate immune response. EMBO Mol Med 7, 547–561 (2015).

13. Osta, B., Benedetti, G. & Miossec, P. Classical and Paradoxical Effects of TNF-alpha on Bone Homeostasis. Front Immunol 5, 48 (2014).

14. Glass, G.E. et al. TNF-alpha promotes fracture repair by augmenting the recruitment and differentiation of muscle-derived stromal cells. Proceedings of the National Academy of Sciences of the United States of America 108, 1585–1590 (2011).

15. Cao, Y., Jansen, I.D.C., Sprangers, S., de Vries, T.J. & Everts, V. TNF-alpha has both stimulatory and inhibitory effects on mouse monocyte-derived osteoclastogenesis. Journal of cellular physiology 232, 3273–3285 (2017).

16. Cho, D.I. et al. Mesenchymal stem cells reciprocally regulate the M1/M2 balance in mouse bone marrow-derived macrophages. Exp Mol Med 46, e70 (2014).

17. Alexander, K.A. et al. Osteal macrophages promote in vivo intramembranous bone healing in a mouse tibial injury model. Journal of bone and mineral research: the official journal of the American Society for Bone and Mineral Research 26, 1517–1532 (2011).

18. Raggatt, L.J. et al. Fracture healing via periosteal callus formation requires macrophages for both initiation and progression of early endochondral ossification. Am J Pathol 184, 3192–3204 (2014).

19. Schlundt, C. et al. Macrophages in bone fracture healing: Their essential role in endochondral ossification. Bone 106, 78–89 (2018).

20. Basanta, D., Gatenby, R.A. & Anderson, A.R. Exploiting evolution to treat drug resistance: combination therapy and the double bind. Mol Pharm 9, 914–921 (2012).

21. Eikenberry, S.E., Nagy, J.D. & Kuang, Y. The evolutionary impact of androgen levels on prostate cancer in a multi-scale mathematical model. Biology direct 5, 24 (2010).

22. Gatenby, R.A., Silva, A.S., Gillies, R.J. & Frieden, B.R. Adaptive therapy. Cancer research 69, 4894–4903 (2009).

23. Horn, M. et al. Model-based decision rules reduce the risk of molecular relapse after cessation of tyrosine kinase inhibitor therapy in chronic myeloid leukemia. Blood 121, 378–384 (2013).

24. Leder, K. et al. Mathematical modeling of PDGF-driven glioblastoma reveals optimized radiation dosing schedules. Cell 156, 603–616 (2014).

25. Rockne, R., Alvord, E.C., Jr., Rockhill, J.K. & Swanson, K.R. A mathematical model for brain tumor response to radiation therapy. J Math Biol 58, 561–578 (2009).

26. Swanson, K.R. et al. Quantifying the role of angiogenesis in malignant progression of gliomas: in silico modeling integrates imaging and histology. Cancer research 71, 7366–7375 (2011).

27. Swanson, K.R., Rostomily, R.C. & Alvord, E.C., Jr. A mathematical modelling tool for predicting survival of individual patients following resection of glioblastoma: a proof of principle. Br J Cancer 98, 113–119 (2008).

28. Anderson, A.R. & Quaranta, V. Integrative mathematical oncology. Nature reviews. Cancer 8, 227–234 (2008).

29. Araujo, A., Cook, L.M., Lynch, C.C. & Basanta, D. An integrated computational model of the bone microenvironment in bone-metastatic prostate cancer. Cancer research 74, 2391–2401 (2014).

30. Komarova, S.V. Mathematical model of paracrine interactions between osteoclasts and osteoblasts predicts anabolic action of parathyroid hormone on bone. Endocrinology 146, 3589–3595 (2005).

31. Pivonka, P. et al. Model structure and control of bone remodeling: a theoretical study. Bone 43, 249–263 (2008).

32. Lemaire, V., Tobin, F.L., Greller, L.D., Cho, C.R. & Suva, L.J. Modeling the interactions between osteoblast and osteoclast activities in bone remodeling. J Theor Biol 229, 293–309 (2004).

33. Ayati, B.P., Edwards, C.M., Webb, G.F. & Wikswo, J.P. A mathematical model of bone remodeling dynamics for normal bone cell populations and myeloma bone disease. Biology direct 5, 28 (2010).

34. Buenzli, P.R., Pivonka, P., Gardiner, B.S. & Smith, D.W. Modelling the anabolic response of bone using a cell population model. J Theor Biol 307, 42–52 (2012).

35. Graham, J.M., Ayati, B.P., Holstein, S.A. & Martin, J.A. The role of osteocytes in targeted bone remodeling: a mathematical model. PLoS One 8, e63884 (2013).

36. Bailon-Plaza, A. & van der Meulen, M.C. A mathematical framework to study the effects of growth factor influences on fracture healing. J Theor Biol 212, 191–209 (2001).

37. Ryser, M.D., Nigam, N. & Komarova, S.V. Mathematical modeling of spatio-temporal dynamics of a single bone multicellular unit. Journal of bone and mineral research: the official journal of the American Society for Bone and Mineral Research 24, 860–870 (2009).

38. Trejo, I.K., Hristo; Chen-Charpentier, Benito Modeling the Macrophage-Mediated Inflammation Involved in the Bone Fracture Healing Process. Mathematical and Computational Applications 24 (2019).

39. Kojouharov, H.V., Trejo, I. & Chen-Charpentier, B.M (2017).

40. Aspenberg, P. & Sandberg, O. Distal radial fractures heal by direct woven bone formation. Acta Orthop 84, 297–300 (2013).

41. Chang, M.K. et al. Osteal tissue macrophages are intercalated throughout human and mouse bone lining tissues and regulate osteoblast function in vitro and in vivo. Journal of immunology (Baltimore, Md.: 1950) 181, 1232–1244 (2008).

42. Sinder, B.P. et al. Bone Mass Is Compromised by the Chemotherapeutic Trabectedin in Association With Effects on Osteoblasts and Macrophage Efferocytosis. Journal of bone and mineral research: the official journal of the American Society for Bone and Mineral Research 32, 2116–2127 (2017).

43. Everts, V. et al. The Bone Lining Cell: Its Role in Cleaning Howship,s Lacunae and Initiating Bone Formation. Journal of Bone and Mineral Research 17, 77–90 (2002).

44. Manolagas, S.C. Birth and death of bone cells: basic regulatory mechanisms and implications for the pathogenesis and treatment of osteoporosis. Endocr Rev 21, 115–137 (2000).

45. McArdle, A. et al. The role and regulation of osteoclasts in normal bone homeostasis and in response to injury. Plastic and reconstructive surgery 135, 808–816 (2015).

46. Raggatt, L.J. & Partridge, N.C. Cellular and molecular mechanisms of bone remodeling. The Journal of biological chemistry 285, 25103–25108 (2010).

47. Hiltunen, A., Vuorio, E. & Aro, H.T. A standardized experimental fracture in the mouse tibia. J Orthop Res 11, 305–312 (1993).

48. Premnath, P. et al. p21(-/-) mice exhibit enhanced bone regeneration after injury. BMC Musculoskelet Disord 18, 435 (2017).

49. Taiani, J.T. et al. Embryonic stem cells incorporate into newly formed bone and do not form tumors in an immunocompetent mouse fracture model. Cell Transplant 22, 1453–1462 (2013).

50. Guihard, P. et al. Oncostatin m, an inflammatory cytokine produced by macrophages, supports intramembranous bone healing in a mouse model of tibia injury. Am J Pathol 185, 765–775 (2015).

51. Vi, L. et al. Macrophages promote osteoblastic differentiation in-vivo: implications in fracture repair and bone homeostasis. Journal of bone and mineral research: the official journal of the American Society for Bone and Mineral Research 30, 1090–1102 (2015).

52. Kon, T. et al. Expression of osteoprotegerin, receptor activator of NF-kappaB ligand (osteoprotegerin ligand) and related proinflammatory cytokines during fracture healing. Journal of bone and mineral research: the official journal of the American Society for Bone and Mineral Research 16, 1004–1014 (2001).

53. Thompson, E.R., Baylink, D.J. & Wergedal, J.E. Increases in number and size of osteoclasts in response to calcium or phosphorus deficiency in the rat. Endocrinology 97, 283–289 (1975).

54. Jeganathan, S., Fiorino, C., Naik, U., Sun, H.s. & Harrison, R.E. Modulation of Osteoclastogenesis with Macrophage M1- and M2-Inducing Stimuli. PLoS ONE 9, e104498 (2014).

55. Zhang, Y.H., Heulsmann, A., Tondravi, M.M., Mukherjee, A. & Abu-Amer, Y. Tumor necrosis factor-alpha (TNF) stimulates RANKL-induced osteoclastogenesis via coupling of TNF type 1 receptor and RANK signaling pathways. The Journal of biological chemistry 276, 563–568 (2001).

56. Zhao, Z. et al. TNF Induction of NF-kappaB RelB Enhances RANKL-Induced Osteoclastogenesis by Promoting Inflammatory Macrophage Differentiation but also Limits It through Suppression of NFATc1 Expression. PLoS One 10, e0135728 (2015).

57. Bendixen, A.C. et al. IL-4 inhibits osteoclast formation through a direct action on osteoclast precursors via peroxisome proliferator-activated receptor gamma 1. Proceedings of the National Academy of Sciences of the United States of America 98, 2443–2448 (2001).

58. Cao, Y. et al. IL-1beta differently stimulates proliferation and multinucleation of distinct mouse bone marrow osteoclast precursor subsets. Journal of leukocyte biology 100, 513–523 (2016).

59. Frost, A. et al. Interleukin (IL)-13 and IL-4 inhibit proliferation and stimulate IL-6 formation in human osteoblasts: evidence for involvement of receptor subunits IL-13R, IL-13Ralpha, and IL-4Ralpha. Bone 28, 268–274 (2001).

60. Kobayashi, K. et al. Tumor necrosis factor alpha stimulates osteoclast differentiation by a mechanism independent of the ODF/RANKL-RANK interaction. J Exp Med 191, 275–286 (2000).

61. Li, X. et al. Parathyroid hormone stimulates osteoblastic expression of MCP-1 to recruit and increase the fusion of pre/osteoclasts. The Journal of biological chemistry 282, 33098–33106 (2007).

62. Nakao, K. et al. IGF2 modulates the microenvironment for osteoclastogenesis. Biochemical and biophysical research communications 378, 462–466 (2009).

63. Song, L. et al. Interleukin-17A facilitates osteoclast differentiation and bone resorption via activation of autophagy in mouse bone marrow macrophages. Mol Med Rep 19, 4743–4752 (2019).

64. Yamada, A. et al. Interleukin-4 inhibition of osteoclast differentiation is stronger than that of interleukin-13 and they are equivalent for induction of osteoprotegerin production from osteoblasts. Immunology 120, 573–579 (2007).

65. Jilka, R.L. The relevance of mouse models for investigating age-related bone loss in humans. J Gerontol A Biol Sci Med Sci 68, 1209–1217 (2013).

66. Kanehisa, J. & Heersche, J.N. Osteoclastic bone resorption: in vitro analysis of the rate of resorption and migration of individual osteoclasts. Bone 9, 73–79 (1988).

67. Gong, L., Zhao, Y., Zhang, Y. & Ruan, Z. The Macrophage Polarization Regulates MSC Osteoblast Differentiation in vitro. Annals of clinical and laboratory science 46, 65–71 (2016).

68. Michalski, M.N., Koh, A.J., Weidner, S., Roca, H. & McCauley, L.K. Modulation of Osteoblastic Cell Efferocytosis by Bone Marrow Macrophages. J Cell Biochem 117, 2697–2706 (2016).

69. Alexander, K.A. et al. Resting and injury-induced inflamed periosteum contain multiple macrophage subsets that are located at sites of bone growth and regeneration. Immunol Cell Biol 95, 7–16 (2017).

70. Orecchioni, M., Ghosheh, Y., Pramod, A.B. & Ley, K. Macrophage Polarization: Different Gene Signatures in M1(LPS+) vs. Classically and M2(LPS–) vs. Alternatively Activated Macrophages. Frontiers in Immunology 10 (2019).

71. Sica, A. & Mantovani, A. Macrophage plasticity and polarization: in vivo veritas. J Clin Invest 122, 787–795 (2012).

72. Mosser, D.M. & Edwards, J.P. Exploring the full spectrum of macrophage activation. Nat Rev Immunol 8, 958–969 (2008).

73. Misharin, A.V., Morales-Nebreda, L., Mutlu, G.M., Budinger, G.R. & Perlman, H. Flow cytometric analysis of macrophages and dendritic cell subsets in the mouse lung. Am J Respir Cell Mol Biol 49, 503–510 (2013).

74. Italiani, P. & Boraschi, D. From Monocytes to M1/M2 Macrophages: Phenotypical vs. Functional Differentiation. Front Immunol 5, 514 (2014).

75. Koh, T.J. & DiPietro, L.A. Inflammation and wound healing: the role of the macrophage. Expert Rev Mol Med 13, e23 (2011).

76. Lo, C.H. & Lynch, C.C. Multifaceted Roles for Macrophages in Prostate Cancer Skeletal Metastasis. Frontiers in endocrinology 9, 247 (2018).

77. Mantovani, A., Biswas, S.K., Galdiero, M.R., Sica, A. & Locati, M. Macrophage plasticity and polarization in tissue repair and remodelling. The Journal of pathology 229, 176–185 (2013).

78. Rose, S., Misharin, A. & Perlman, H. A novel Ly6C/Ly6G-based strategy to analyze the mouse splenic myeloid compartment. Cytometry A 81, 343–350 (2012).

79. Macrophage activation unveiled. J Exp Med 202, 884 (2005).

80. Liyanage, S.E. et al. Flow cytometric analysis of inflammatory and resident myeloid populations in mouse ocular inflammatory models. Exp Eye Res 151, 160–170 (2016).

81. Minutti, C.M., Knipper, J.A., Allen, J.E. & Zaiss, D.M.W. Tissue-specific contribution of macrophages to wound healing. Seminars in cell & developmental biology 61, 3–11 (2017).

82. Boniakowski, A.E., Kimball, A.S., Jacobs, B.N., Kunkel, S.L. & Gallagher, K.A. Macrophage-Mediated Inflammation in Normal and Diabetic Wound Healing. Journal of immunology (Baltimore, Md.: 1950) 199, 17–24 (2017).

83. Yakubenko, V.P. et al. Oxidative modifications of extracellular matrix promote the second wave of inflammation via beta2 integrins. Blood 132, 78–88 (2018).

84. Batoon, L., Millard, S.M., Raggatt, L.J. & Pettit, A.R. Osteomacs and Bone Regeneration. Current osteoporosis reports 15, 385–395 (2017).

85. Batoon, L. et al. CD169(+) macrophages are critical for osteoblast maintenance and promote intramembranous and endochondral ossification during bone repair. Biomaterials (2017).

86. Bozec, A. & Soulat, D. Latest perspectives on macrophages in bone homeostasis. Pflügers Archiv - European Journal of Physiology 469, 517–525 (2017).

87. Charles, J.F. et al. Inflammatory arthritis increases mouse osteoclast precursors with myeloid suppressor function. The Journal of Clinical Investigation 122, 4592–4605.

88. Estus, T.L., Choudhary, S. & Pilbeam, C.C. Prostaglandin-mediated inhibition of PTH-stimulated β-catenin signaling in osteoblasts by bone marrow macrophages. Bone 85, 123–130 (2016).

89. Guihard, P. et al. Induction of Osteogenesis in Mesenchymal Stem Cells by Activated Monocytes/Macrophages Depends on Oncostatin M Signaling. STEM CELLS 30, 762–772 (2012).

90. He, D. et al. M1-like Macrophage Polarization Promotes Orthodontic Tooth Movement. Journal of dental research 94, 1286–1294 (2015).

91. Huang, R., Wang, X., Zhou, Y. & Xiao, Y. RANKL-induced M1 macrophages are involved in bone formation. Bone research 5, 17019 (2017).

92. Lam, J. et al. TNF-alpha induces osteoclastogenesis by direct stimulation of macrophages exposed to permissive levels of RANK ligand. J Clin Invest 106, 1481–1488 (2000).

93. Yamaguchi, T. et al. Proinflammatory M1 Macrophages Inhibit RANKL-Induced Osteoclastogenesis. Infection and immunity 84, 2802–2812 (2016).

94. Kaur, S. et al. Role of bone marrow macrophages in controlling homeostasis and repair in bone and bone marrow niches. Seminars in cell & developmental biology 61, 12–21 (2017).

95. Sinder, B.P., Pettit, A.R. & McCauley, L.K. Macrophages: Their Emerging Roles in Bone. Journal of Bone and Mineral Research 30, 2140–2149 (2015).

96. Davis, M.J. et al. Macrophage M1/M2 polarization dynamically adapts to changes in cytokine microenvironments in Cryptococcus neoformans infection. mBio 4, e00264–00213 (2013).

97. Michlewska, S., Dransfield, I., Megson, I.L. & Rossi, A.G. Macrophage phagocytosis of apoptotic neutrophils is critically regulated by the opposing actions of pro-inflammatory and anti-inflammatory agents: key role for TNF-alpha. FASEB journal: official publication of the Federation of American Societies for Experimental Biology 23, 844–854 (2009).

98. Crane, M.J. et al. The monocyte to macrophage transition in the murine sterile wound. PLoS One 9, e86660 (2014).

99. Lampiasi, N., Russo, R. & Zito, F. The Alternative Faces of Macrophage Generate Osteoclasts. Biomed Res Int 2016, 9089610 (2016).

100. Das, A., Ganesh, K., Khanna, S., Sen, C.K. & Roy, S. Engulfment of apoptotic cells by macrophages: a role of microRNA-21 in the resolution of wound inflammation. Journal of immunology (Baltimore, Md.: 1950) 192, 1120–1129 (2014).

101. Lu, G. et al. Myeloid cell-derived inducible nitric oxide synthase suppresses M1 macrophage polarization. Nature communications 6, 6676 (2015).

