## Supplemental Material for "Computational modeling reveals a key role for polarized myeloid cells in controlling osteoclast activity during bone injury repair"

**AFFILIATIONS:**
[1]Tumor Biology Department and [2]Integrated Mathematical Oncology Department, Moffitt Cancer Center and Research Institute, Tampa, Florida.
[3]Oncology and Cancer Biology PhD Program, University of South Florida, Tampa, Florida.

[#]Co-First Author
[*]Co-Senior Author

**CORRESPONDING AUTHORS**: Conor C Lynch and David Basanta, SRB, Moffitt Cancer Center, 12902 Magnolia Drive, Tampa, FL, USA. Phone: 813-745-8094. Fax: 813-745-3829.

*Lo & Baratchart et al.* ***Supplemental Fig. 1***

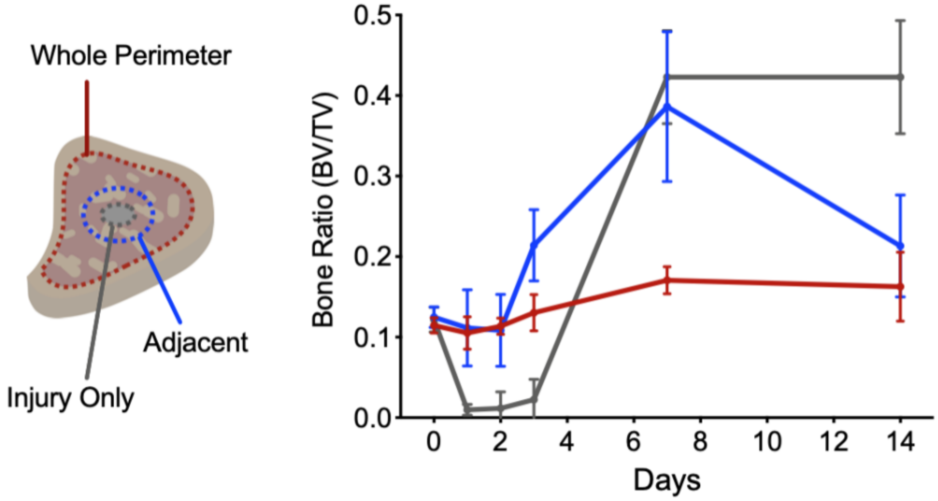


**Supplemental Fig. 1.** Analyses of bone status reveal most intense bone remodeling at site of bone injury. Micro-computed tomography analysis was performed using scanned data from three sets of contours for each bone: 1) the whole trabecular perimeter (Red), 2) the injury site and adjacent tissue (Blue) and 3) the injury only (Grey). Data is plotted as total bone volume over total volume (BV/TV).

*Lo & Baratchart et al.* ***Supplemental Fig. 2***

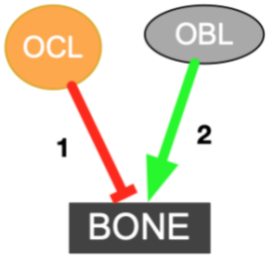


| Description | Supplemental References |
|---|---|
| Bone is maintained at homeostasis by osteoclast and osteoblast | *1-9* |
| Bone undergoes heightened remodeling during injury repair (**1**, **2**) | *10-19* |
| Osteoclastic bone resorption (**1**) | *2, 7, 8, 14, 20-28* |
| Osteoblastic bone formation (**2**) | *6, 14-16, 19, 23, 24, 29, 30* |

**Supplemental Fig. 2.** Relationship between osteoblast, osteoclast and bone per published literature. Number of publications were consulted not only to derive relationship between three species but also numerical parameters applicable for ODE modeling.

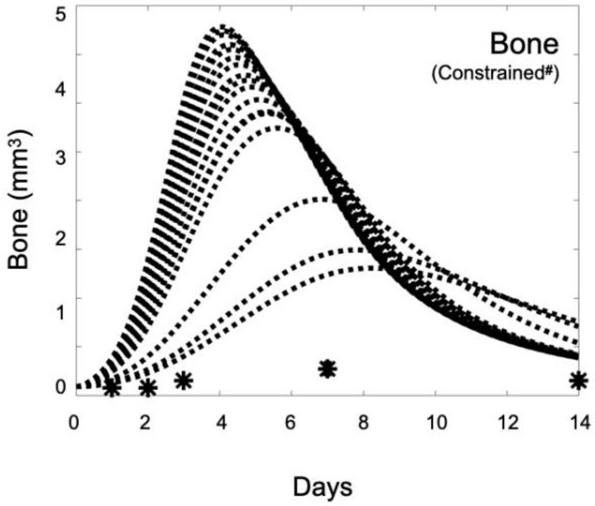


**Supplemental Fig. 3.** Testing fits to bone dynamic using published constant osteoclast resorption rates. Representative model iterations testing bone resorption rates equally spaced across the literature-derived range of possible values.

*Lo & Baratchart et al.* ***Supplemental Fig. 4***

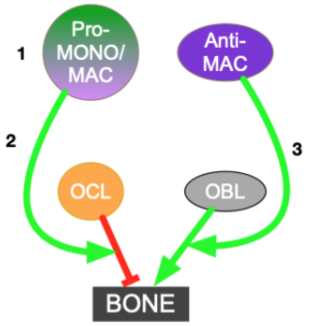


| Description | Supplemental References |
|---|---|
| Myeloid cells polarize to M1 and M2 states during wounding and repair (1) | *3, 29, 30, 35-45* |
| Pro-inflammatory myeloid cells stimulate osteoclast bone resorption (2) | *24, 29, 42, 44, 46-55* |
| Anti-inflammatory myeloid cells stimulate osteoblast bone formation (3) | *29, 36, 41, 42, 45, 49, 50, 52-63* |

**Supplemental Fig. 4.** Relationship between pro- and anti-inflammatory myeloid cells and osteoblast, osteoclast and bone. Numerous published literatures were consulted to not only derive relationship between myeloid and bone cells, but also to derive quantitative parameters potentially useful in ODE modeling.

*Lo & Baratchart et al.* ***Supplemental Fig. 5***

**a**

| Population | Markers |
|---|---|
| MONO | CD11b$^+$ LY-6C$^{HI}$ LY-6G$^-$ |
| MAC | CD11b$^+$ LY-6C$^{LO}$ LY-6G$^-$ |
| Pro-MONO | CD11b$^+$ LY-6C$^{HI}$ LY-6G$^-$ NOS2$^+$ |
| Anti-MONO | CD11b$^+$ LY-6C$^{HI}$ LY-6G$^-$ ARG1$^+$ |
| Pro-MAC | CD11b$^+$ LY-6C$^{LO}$ LY-6G$^-$ NOS2$^+$ |
| Anti-MAC | CD11b$^+$ LY-6C$^{LO}$ LY-6G$^-$ ARG1$^+$ |

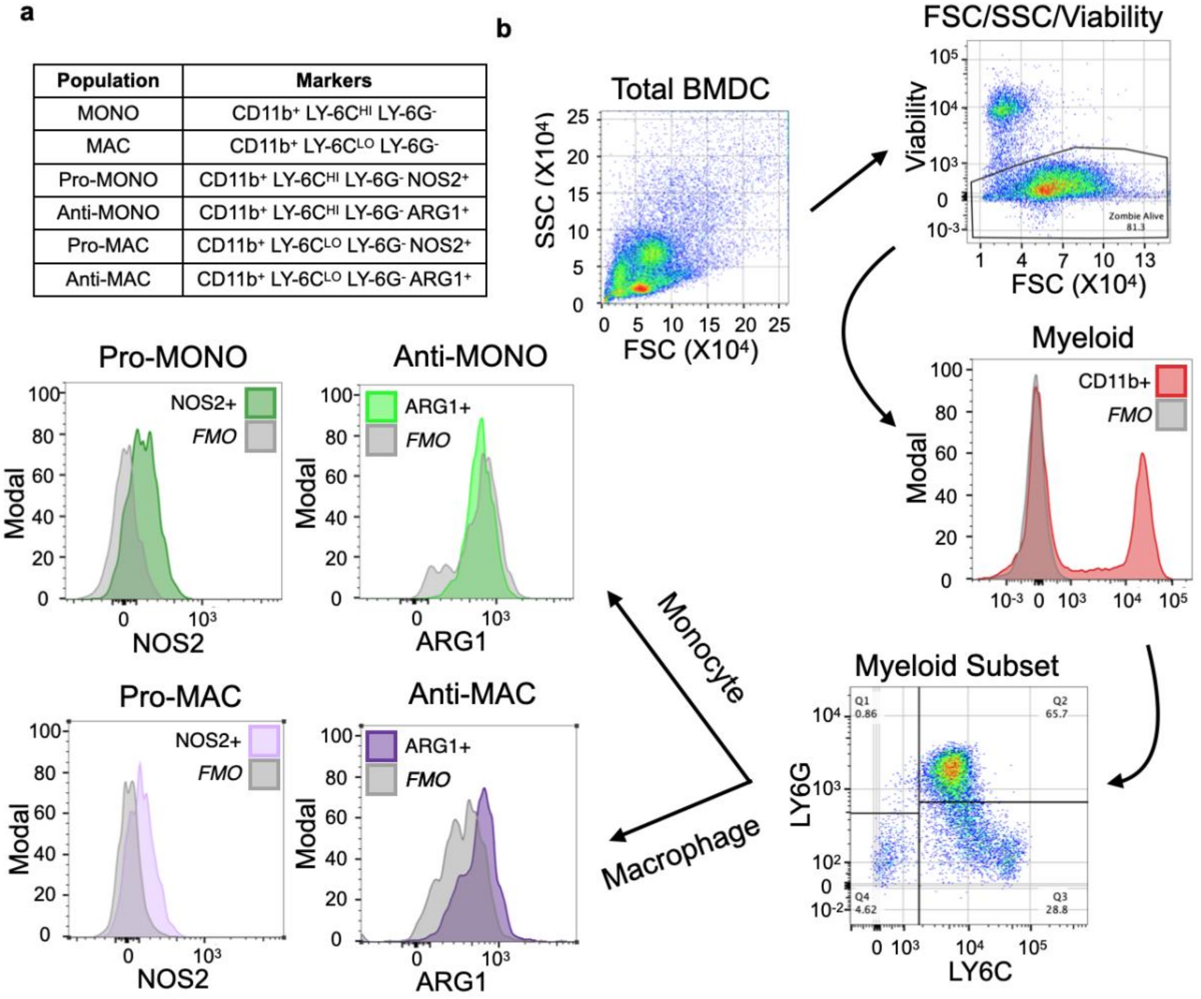


**Supplemental Fig. 5.** Flow cytometry gating strategy for detection of polarized monocytes and macrophages. **a** table detailing combination of markers used to discern each subpopulation of interest within the heterogeneous bone marrow. **b** flow chart detailing the process by which the whole bone marrow is gradually dissected to reveal the cells of interest.

*Lo & Baratchart et al.* ***Supplemental Fig. 6***

**a**

$$\frac{\mathrm{d}OB}{\mathrm{d}t} = H_{OB} + \frac{\gamma_{OB}OB(Tanab - t)(t < Tanab)}{Tanab} - \delta_{OB}OB$$

$$\frac{\mathrm{d}OC}{\mathrm{d}t} = -Inhib_{OC}OC(t < TantiCatab) + R_{OC}(t > Tcatab) - R_{OC}OC/OC_0$$

$$\frac{\mathrm{d}B}{\mathrm{d}t} = -\delta_B(t)OCB + \Pi_B OB$$

$$\delta_B(t) = (\delta_{B_0} + \delta_{B_1}t)(t < 1) + (\delta_{B_0} + \delta_{B_1} + \delta_{B_2}(t-1))(t \geq 1)(t < 2)$$
$$+ (\delta_{B_0} + \delta_{B_1} + \delta_{B_2} + \delta_{B_3}(t-2))(t \geq 2)(t < 3)$$
$$+ (\delta_{B_0} + \delta_{B_1} + \delta_{B_2} + \delta_{B_3} + \delta_{B_4}(t-3))(t \geq 3)(t < 7)$$
$$+ (\delta_{B_0} + \delta_{B_1} + \delta_{B_2} + \delta_{B_3} + 4\delta_{B_4} + (t-7)\delta_{B_0})(t \geq 7)$$

**b**

**c**

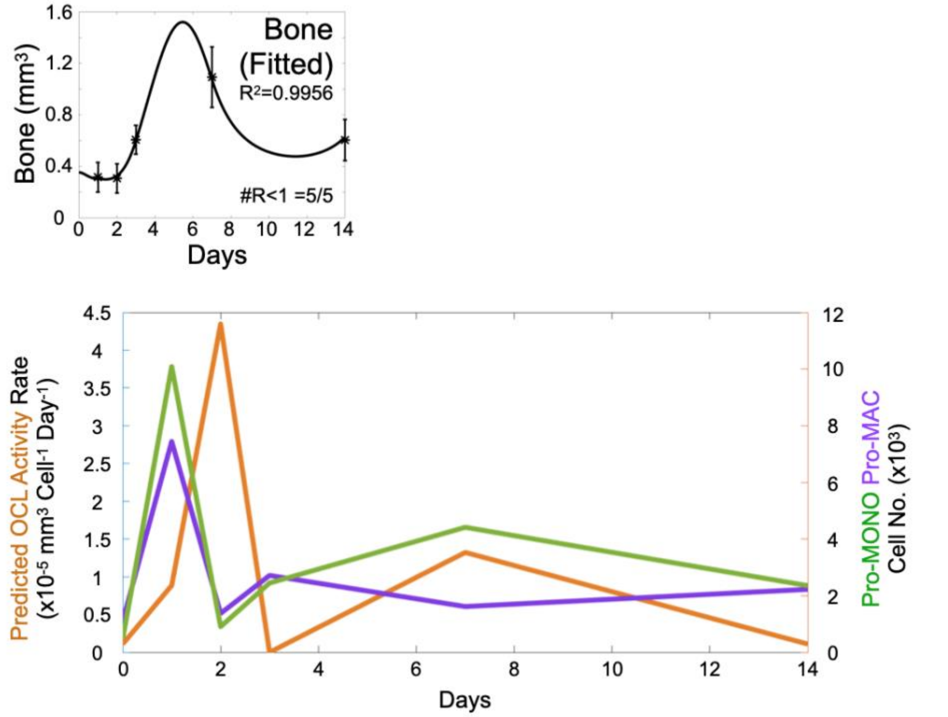


**Supplemental Fig. 6.** Agnostic approach showing necessary temporal changes to osteoclast activity to recapitulate bone dynamics. **a** constraint on osteoclast resorption was loosened in initial osteoblast-osteoclast-bone model to ask what are temporal changes necessary to allow accurate recapitulation of bone dynamics. **b** model was able to generate accurate bone fits given this loosened constraint. **c** predicted osteoclast activity dynamics was plotted against experimental pro-inflammatory monocyte and macrophage dynamics to demonstrate similarity.

*Lo & Baratchart et al.* ***Supplemental Fig. 7***

**a**

$$\frac{\mathrm{d}OB}{\mathrm{d}t} = H_{OB} + \frac{\gamma_{OB} OB(Tanab - t)(t < Tanab)}{Tanab} - \delta_{OB} OB$$

$$\frac{\mathrm{d}OC}{\mathrm{d}t} = -Inhib_{OC} OC(t < TantiCatab) + R_{OC}(t > Tcatab) - R_{OC} OC/OC_0$$

$$\frac{\mathrm{d}B}{\mathrm{d}t} = -\delta_B OCB + \Pi_B(t) OB$$

$$\Pi_B(t) = (\Pi_{B_0} + \Pi_{B_1} t)(t < 1) + (\Pi_{B_0} + \Pi_{B_1} + \Pi_{B_2}(t-1))(t \geq 1)(t < 2)$$
$$+ (\Pi_{B_0} + \Pi_{B_1} + \Pi_{B_2} + \Pi_{B_3}(t-2))(t \geq 2)(t < 3)$$
$$+ (\Pi_{B_0} + \Pi_{B_1} + \Pi_{B_2} + \Pi_{B_3} + \Pi_{B_4}(t-3))(t \geq 3)(t < 7)$$
$$+ (\Pi_{B_0} + \Pi_{B_1} + \Pi_{B_2} + \Pi_{B_3} + 4\Pi_{B_4} + (t-7)\Pi_{B_0})(t \geq 7)$$

**b**

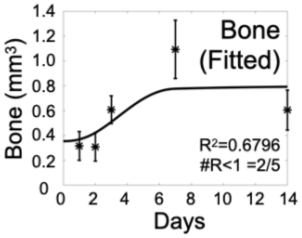


**Supplemental Fig. 7.** Agnostic approach showing temporal changes to osteoblast activity does not recapitulate bone dynamics. **a** constraint to osteoblast resorption was loosened in initial osteoblast-osteoclast-bone model to ask what are temporal changes necessary to allow accurate recapitulation of bone dynamics. **b** even with the loosened constraint, model failed to generate plausible bone fits.

**Supplemental References.** A number of additional references were consulted to characterize and parameterize biological behaviors and relationships between cellular species.


1. Lampiasi, N., Russo, R. & Zito, F. The Alternative Faces of Macrophage Generate Osteoclasts. BioMed Research International 2016, 9 (2016).
2. Asagiri, M. & Takayanagi, H. The molecular understanding of osteoclast differentiation. Bone 40, 251-264 (2007).
3. Chang, C.F. et al. Erythrocyte efferocytosis modulates macrophages towards recovery after intracerebral hemorrhage. J Clin Invest 128, 607-624 (2018).
4. Rodan, G.A. Bone homeostasis. Proceedings of the National Academy of Sciences of the United States of America 95, 13361-13362 (1998).
5. Zaidi, M., Yuen, T., Sun, L. & Rosen, C.J. Regulation of Skeletal Homeostasis. Endocrine Reviews 39, 701-718 (2018).
6. Everts, V. et al. The bone lining cell: its role in cleaning Howship's lacunae and initiating bone formation. Journal of bone and mineral research : the official journal of the American Society for Bone and Mineral Research 17, 77-90 (2002).
7. Li, X. et al. Parathyroid hormone stimulates osteoblastic expression of MCP-1 to recruit and increase the fusion of pre/osteoclasts. The Journal of biological chemistry 282, 33098-33106 (2007).
8. McArdle, A. et al. The role and regulation of osteoclasts in normal bone homeostasis and in response to injury. Plastic and reconstructive surgery 135, 808-816 (2015).
9. Raggatt, L.J. & Partridge, N.C. Cellular and Molecular Mechanisms of Bone Remodeling. Journal of Biological Chemistry 285, 25103-25108 (2010).
10. Wang, L. et al. Osteoblast-induced osteoclast apoptosis by fas ligand/FAS pathway is required for maintenance of bone mass. Cell Death Differ 22, 1654-1664 (2015).
11. Nijweide, P.J., Burger, E.H. & Feyen, J.H. Cells of bone: proliferation, differentiation, and hormonal regulation. Physiol Rev 66, 855-886 (1986).
12. Pakyari, M., Farrokhi, A., Maharlooei, M.K. & Ghahary, A. Critical Role of Transforming Growth Factor Beta in Different Phases of Wound Healing. Advances in wound care 2, 215-224 (2013).
13. Grundnes, O. & Reikeras, O. The importance of the hematoma for fracture healing in rats. Acta orthopaedica Scandinavica 64, 340-342 (1993).
14. Campbell, T.M., Wong, W.T. & Mackie, E.J. Establishment of a model of cortical bone repair in mice. Calcif Tissue Int 73, 49-55 (2003).
15. Loi, F. et al. Inflammation, fracture and bone repair. Bone 86, 119-130 (2016).
16. Schindeler, A., McDonald, M.M., Bokko, P. & Little, D.G. Bone remodeling during fracture repair: The cellular picture. Semin Cell Dev Biol 19, 459-466 (2008).
17. Claes, L., Recknagel, S. & Ignatius, A. Fracture healing under healthy and inflammatory conditions. Nature reviews. Rheumatology 8, 133-143 (2012).
18. Sims, N.A. & Martin, T.J. Coupling the activities of bone formation and resorption: a multitude of signals within the basic multicellular unit. BoneKEy reports 3, 481 (2014).
19. Raggatt, L.J. & Partridge, N.C. Cellular and molecular mechanisms of bone remodeling. The Journal of biological chemistry 285, 25103-25108 (2010).
20. Takahashi, M. et al. Docetaxel inhibits bone resorption through suppression of osteoclast formation and function in different manners. Journal of bone and mineral metabolism 27, 24-35 (2009).
21. Teitelbaum, S.L. Bone resorption by osteoclasts. Science 289, 1504-1508 (2000).
22. Yavropoulou, M.P. & Yovos, J.G. Osteoclastogenesis--current knowledge and future perspectives. Journal of musculoskeletal & neuronal interactions 8, 204-216 (2008).

23. Manolagas, S.C. Birth and death of bone cells: basic regulatory mechanisms and implications for the pathogenesis and treatment of osteoporosis. Endocr Rev 21, 115-137 (2000).
24. Levaot, N. et al. Osteoclast fusion is initiated by a small subset of RANKL-stimulated monocyte progenitors, which can fuse to RANKL-unstimulated progenitors. Bone 79, 21-28 (2015).
25. Aubin, J.E. & Bonnelye, E. Osteoprotegerin and its ligand: a new paradigm for regulation of osteoclastogenesis and bone resorption. Osteoporos Int 11, 905-913 (2000).
26. Jilka, R.L. The relevance of mouse models for investigating age-related bone loss in humans. J Gerontol A Biol Sci Med Sci 68, 1209-1217 (2013).
27. Kanehisa, J. & Heersche, J.N. Osteoclastic bone resorption: in vitro analysis of the rate of resorption and migration of individual osteoclasts. Bone 9, 73-79 (1988).
28. Thompson, E.R., Baylink, D.J. & Wergedal, J.E. Increases in number and size of osteoclasts in response to calcium or phosphorus deficiency in the rat. Endocrinology 97, 283-289 (1975).
29. Wu, A.C., Raggatt, L.J., Alexander, K.A. & Pettit, A.R. Unraveling macrophage contributions to bone repair. Bonekey Rep 2, 373 (2013).
30. Guihard, P. et al. Induction of osteogenesis in mesenchymal stem cells by activated monocytes/macrophages depends on oncostatin M signaling. Stem Cells 30, 762-772 (2012).
31. Wei, W. et al. Osteoclast progenitors reside in the peroxisome proliferator-activated receptor gamma-expressing bone marrow cell population. Mol Cell Biol 31, 4692-4705 (2011).
32. Akchurin, T. et al. Complex dynamics of osteoclast formation and death in long-term cultures. PLoS One 3, e2104 (2008).
33. Proctor, C.J. & Gartland, A. Simulated Interventions to Ameliorate Age-Related Bone Loss Indicate the Importance of Timing. Front Endocrinol (Lausanne) 7, 61 (2016).
34. Jilka, R.L., Weinstein, R.S., Bellido, T., Parfitt, A.M. & Manolagas, S.C. Osteoblast programmed cell death (apoptosis): modulation by growth factors and cytokines. J Bone Miner Res 13, 793-802 (1998).
35. de Bruin, A.M. et al. IFNγ induces monopoiesis and inhibits neutrophil development during inflammation. Blood 119, 1543-1554 (2012).
36. Mantovani, A., Biswas, S.K., Galdiero, M.R., Sica, A. & Locati, M. Macrophage plasticity and polarization in tissue repair and remodelling. The Journal of pathology 229, 176-185 (2013).
37. Zhou, D. et al. Macrophage polarization and function with emphasis on the evolving roles of coordinated regulation of cellular signaling pathways. Cellular signalling 26, 192-197 (2014).
38. Allen, J.E. & Ruckerl, D. The Silent Undertakers: Macrophages Programmed for Efferocytosis. Immunity 47, 810-812 (2017).
39. Liu, Y.C., Zou, X.B., Chai, Y.F. & Yao, Y.M. Macrophage polarization in inflammatory diseases. International journal of biological sciences 10, 520-529 (2014).
40. Michlewska, S., Dransfield, I., Megson, I.L. & Rossi, A.G. Macrophage phagocytosis of apoptotic neutrophils is critically regulated by the opposing actions of pro-inflammatory and anti-inflammatory agents: key role for TNF-alpha. FASEB journal : official publication of the Federation of American Societies for Experimental Biology 23, 844-854 (2009).
41. Michalski, M.N., Koh, A.J., Weidner, S., Roca, H. & McCauley, L.K. Modulation of Osteoblastic Cell Efferocytosis by Bone Marrow Macrophages. J Cell Biochem 117, 2697-2706 (2016).
42. Cho, S.W. Role of osteal macrophages in bone metabolism. J Pathol Transl Med 49, 102-104 (2015).
43. GALEA, E. & FEINSTEIN, D.L. Regulation of the expression of the inflammatory nitric oxide synthase (NOS2) by cyclic AMP. The FASEB Journal 13, 2125-2137 (1999).
44. Pollard, J.W. Trophic macrophages in development and disease. Nat Rev Immunol 9, 259-270 (2009).

45. Cho, S.W. et al. Osteal macrophages support physiologic skeletal remodeling and anabolic actions of parathyroid hormone in bone. Proceedings of the National Academy of Sciences of the United States of America 111, 1545-1550 (2014).
46. Osta, B., Benedetti, G. & Miossec, P. Classical and Paradoxical Effects of TNF-alpha on Bone Homeostasis. Front Immunol 5, 48 (2014).
47. Zhang, Y.H., Heulsmann, A., Tondravi, M.M., Mukherjee, A. & Abu-Amer, Y. Tumor necrosis factor-alpha (TNF) stimulates RANKL-induced osteoclastogenesis via coupling of TNF type 1 receptor and RANK signaling pathways. The Journal of biological chemistry 276, 563-568 (2001).
48. Hurst, S.M. et al. Il-6 and its soluble receptor orchestrate a temporal switch in the pattern of leukocyte recruitment seen during acute inflammation. Immunity 14, 705-714 (2001).
49. Alexander, K.A. et al. Osteal macrophages promote in vivo intramembranous bone healing in a mouse tibial injury model. Journal of bone and mineral research : the official journal of the American Society for Bone and Mineral Research 26, 1517-1532 (2011).
50. Pettit, A.R., Chang, M.K., Hume, D.A. & Raggatt, L.J. Osteal macrophages: a new twist on coupling during bone dynamics. Bone 43, 976-982 (2008).
51. He, D. et al. M1-like Macrophage Polarization Promotes Orthodontic Tooth Movement. Journal of dental research 94, 1286-1294 (2015).
52. Jeganathan, S., Fiorino, C., Naik, U., Sun, H.S. & Harrison, R.E. Modulation of osteoclastogenesis with macrophage M1- and M2-inducing stimuli. PLoS One 9, e104498 (2014).
53. Zhao, Z. et al. TNF Induction of NF-kappaB RelB Enhances RANKL-Induced Osteoclastogenesis by Promoting Inflammatory Macrophage Differentiation but also Limits It through Suppression of NFATc1 Expression. PLoS One 10, e0135728 (2015).
54. Rajfer, R.A. et al. Prevention of Osteoporosis in the Ovariectomized Rat by Oral Administration of a Nutraceutical Combination That Stimulates Nitric Oxide Production. J Osteoporos 2019, 1592328 (2019).
55. Horwood, N.J. Macrophage Polarization and Bone Formation: A review. Clin Rev Allergy Immunol 51, 79-86 (2016).
56. Yamada, A. et al. Interleukin-4 inhibition of osteoclast differentiation is stronger than that of interleukin-13 and they are equivalent for induction of osteoprotegerin production from osteoblasts. Immunology 120, 573-579 (2007).
57. Sinder, B.P. et al. Bone Mass Is Compromised by the Chemotherapeutic Trabectedin in Association With Effects on Osteoblasts and Macrophage Efferocytosis. Journal of bone and mineral research : the official journal of the American Society for Bone and Mineral Research 32, 2116-2127 (2017).
58. Guihard, P. et al. Oncostatin M, an Inflammatory Cytokine Produced by Macrophages, Supports Intramembranous Bone Healing in a Mouse Model of Tibia Injury. The American Journal of Pathology 185, 765-775 (2015).
59. Bozec, A. & Soulat, D. Latest perspectives on macrophages in bone homeostasis. Pflugers Archiv : European journal of physiology 469, 517-525 (2017).
60. Raggatt, L.J. et al. Fracture healing via periosteal callus formation requires macrophages for both initiation and progression of early endochondral ossification. Am J Pathol 184, 3192-3204 (2014).
61. Schlundt, C. et al. Macrophages in bone fracture healing: Their essential role in endochondral ossification. Bone 106, 78-89 (2018).
62. Gong, L., Zhao, Y., Zhang, Y. & Ruan, Z. The Macrophage Polarization Regulates MSC Osteoblast Differentiation in vitro. Annals of clinical and laboratory science 46, 65-71 (2016).
63. Chang, M.K. et al. Osteal tissue macrophages are intercalated throughout human and mouse bone lining tissues and regulate osteoblast function in vitro and in vivo. J Immunol 181, 1232-1244 (2008).

64. Yang, J., Zhang, L., Yu, C., Yang, X.F. & Wang, H. Monocyte and macrophage differentiation: circulation inflammatory monocyte as biomarker for inflammatory diseases. Biomark Res 2, 1 (2014).
65. Italiani, P. & Boraschi, D. From Monocytes to M1/M2 Macrophages: Phenotypical vs. Functional Differentiation. Front Immunol 5, 514 (2014).
